# Non-invasive in vivo acoustoelectric neuromodulation and its contribution to ultrasound stimulation

**DOI:** 10.1101/2025.11.02.685597

**Authors:** Jean L. Rintoul, Christopher Butler, Robin O. Cleveland, Nir Grossman

## Abstract

Non-invasive brain stimulation offers therapeutic potential without the risks of surgery, yet current electrical approaches lack spatial precision and depth due to the long wavelengths of electric fields. Here we demonstrate that the acoustoelectric interaction—the nonlinear coupling between applied acoustic and electric fields—can overcome these limitations to achieve spatially focused, non-invasive neuromodulation. Using *in vitro* and in vivo rodent electrophysiology, we show motor-evoked potentials that depend on both the amplitude and frequency of the acoustoelectric field, with artefactual controls excluding purely acoustic or electrical origins. We further identify an acoustoelectric contribution to conventional ultrasound stimulation, arising from the interaction between ultrasound-induced electrical signals and propagating acoustic waves. These findings establish acoustoelectric neuromodulation as a distinct mechanism of neural activation and a significant contributor to how ultrasound stimulation influences brain activity, opening new directions for precise, non-invasive neuromodulation and neurotherapeutic development.

## INTRODUCTION

Non-invasive neuromodulation requires new approaches to achieve its translational potential. Electrical modalities - including electroconvulsive therapy (ECT)^1^, transcranial magnetic stimulation (TMS)^2^, transcranial electrical stimulation (TES)^3^, and temporal interference (TI) ^4,5^, span a continuum from established practice to rapidly advancing translational innovations, yet all are limited in spatial focality by the long wavelengths of electric fields and their uneven distribution in the brain’s heterogeneous dielectric structure.

In contrast, transcranial focused ultrasound stimulation (tFUS)^6^ offers superior spatial precision among non-invasive modalities, yet the mechanism by which acoustic fields modulate neuronal activity remains incompletely understood. Clarifying this mechanism is essential for improving specificity and safety. Proposed explanations include activation of mechanosensitive ion channels^7^, modulation of microtubules^8^, thermally mediated effects^9^, indirect auditory pathways^10^, cavitation^11^ and the acoustic radiation force^12^. By comparison, electric-field–driven modulation of neuronal excitability is mechanistically described within cable theory and Hodgkin–Huxley^13^ models. Ideally, a neuromodulation technique would be non-invasive, safe, elicit predictable responses at relevant amplitudes, and be selectively targetable to cortical and subcortical regions.

The acoustoelectric effect—the interaction of acoustic and electric fields in ionic media such as saline or brain tissue—may enable electrically mediated neuromodulation with the spatial precision of ultrasound. First described in 1946 by Fox, Herzfeld, and Rock^14^, the phenomenon arises from conductivity changes produced by compression and rarefaction of ions at the acoustic focus. When an electric field is present, these local conductivity fluctuations induce voltage gradients at the ultrasound focal zone. This principle underlies ultrasound current-source-density imaging^15,16^, enabling spatially focused detection of current distributions. We have shown that the acoustoelectric effect is multiplicative (heterodyning)^17^, generating sum- and difference-frequency electric components confined to the acoustic focus.

If the difference frequency falls within electrophysiologically relevant frequencies and amplitudes, the resulting focused electric field could drive neuronal activation via a direct electric mechanism while preserving ultrasound’s focality (**Fig. 1**). This approach offers two independent control axes—acoustic and electric—for optimising amplitude and safety and permits ex vivo calibration by measuring the spatial profile and magnitude of the induced field.

**Figure 1.**
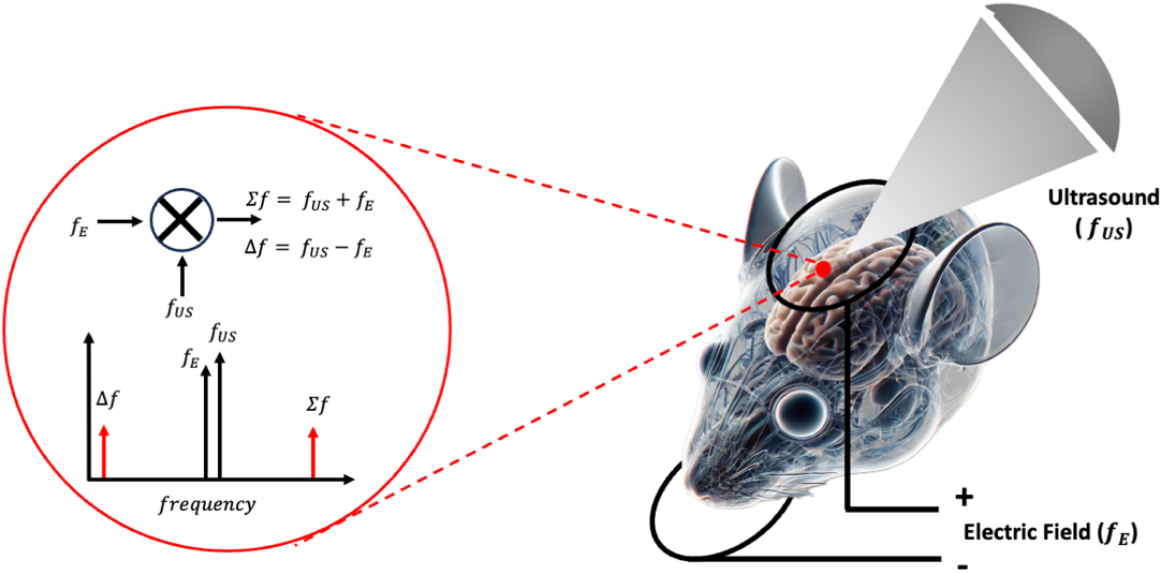
Acoustoelectric neuromodulation and its contribution to ultrasound neuromodulation. **a**, A focal acoustic field *P*(*f*_US_) and a non-focal electric field *E*(*f*_E_) mix via the acoustoelectric interaction, producing electrical signals at the difference frequency *f*_US_ - *f*_E_ and at the sum frequency *f*_US_ + *f*_E_. If the difference frequency Δ*f* is in the range to induce electrophysiological responses, then neuromodulation is possible.

In parallel, the voltage waveform that powers the ultrasound transducer can capacitively couple to surrounding ionic media (e.g., saline or brain tissue), generating secondary electrical artefacts at the acoustic frequency^18–20,^. This coupling is enhanced when a water-filled coupling cone is positioned near the brain. We hypothesised that such artefactual fields can also participate in acoustoelectric mixing, producing a low-frequency component that contributes to tFUS-evoked responses.

Here we test whether acoustoelectrically generated electric fields modulate neural activity *in vitro* (saline phantoms) and *in vivo* (rodents). We find that evoked responses depend on both the amplitude and frequency of the acoustoelectric field and require the concurrent presence of acoustic and electric fields. Having established that acoustoelectric neuromodulation is genuine rather than artefactual, we further show that mixing between the transducer-driven artefactual field and the propagating acoustic wave is a significant contributor to ultrasound-induced brain stimulation. These findings identify the acoustoelectric effect as a key mechanism in ultrasound neuromodulation and support the development of safer, more precise, and mechanistically grounded non-invasive brain-stimulation therapies.

## RESULTS

### Acoustoelectric amplitude characterization

First, we confirmed the focality of the acoustoelectrically generated electrical difference frequency in a tank filled with 0.9% saline (**Fig. 2 a**), by using electrodes to generate an electric field and a 500kHz focused transducer to create a focused ultrasound field at 1MPa. A spatial map of the acoustoelectric field at Δ*f* was measured by translating the stimulating electrode pair and measurement electrode through the acoustic field with 0.5mm steps. It can be seen that the acoustoelectric field in the lateral plane 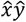 (**Fig. 2 b**) and axial plane 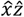 (**Fig. 2 c**) is consistent with the focal region (**Supplemental Note 1**).

**Figure 2.**
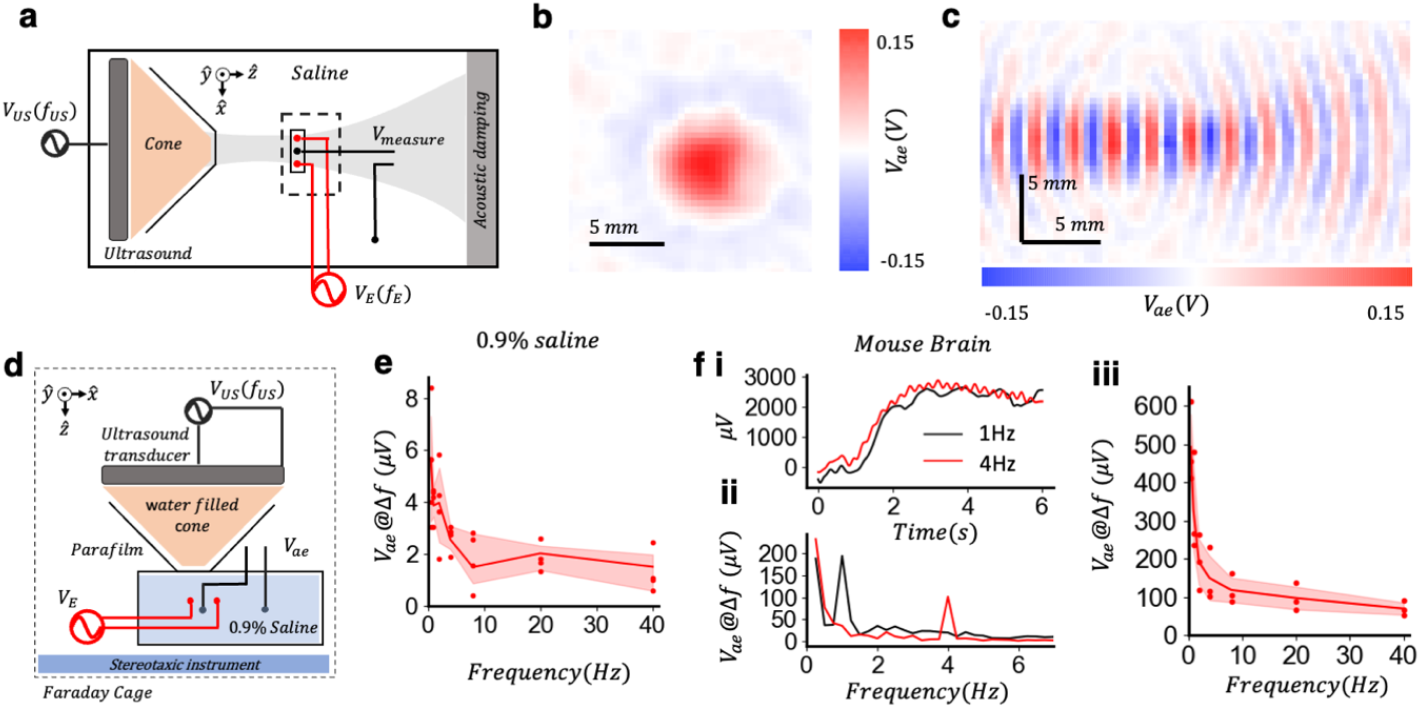
Amplitude characterisation of the acoustoelectric signal. **a**, Free-field phantom measurements using a tank filled with 0.9% saline, a focused ultrasound transducer (500 kHz) generating a focal peak 1MPa, two electrodes separated by 7 mm generating an electric field, and a measurement electrode. **b**, 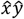 spatial map of acoustoelectric field (for *V*_*E*_= 12V and *f*_*E*_ =8kHz) band-filtered at Δ*f*=492 kHz. **c**, 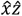 spatial maps of acoustoelectric signal at Δ*f* as in b. **d**, Setup using a petri dish with 0.9% saline to mimic the small volume of a mouse head, within the electrophysiology set up. **e**, Measurements of *V*_*AE*_ as a function of difference frequency (for *V*_*E*_=10V) with 4 measurements at each frequency. **f**, *In vivo* mouse brain measurements (n=3 mice) using an electrode arrangement and settings as in e. i) Representative time series acoustoelectric amplitudes for Δ*f*=1Hz and Δ*f*=4Hz. ii) Amplitude spectrum of the acoustoelectric signal for Δ*f*=1Hz and Δ*f*=4Hz iii) *In vivo* acoustoelectric amplitudes as function of Δ*f*.

The acoustoelectric interaction has previously been reported with very small amplitudes^17,21^ that would be sub-threshold to neural responses^22^. However, it has primarily been explored at higher frequencies close to the transducer frequency^23^, leaving a knowledge gap at the low frequencies (0-100Hz) known to stimulate neurons^24^. To understand acoustoelectric amplitudes at low frequencies, we measured the acoustoelectric voltage amplitudes over a range of low frequencies in a saline filled petri-dish within the intended *in vivo* electrophysiology apparatus (**Fig. 2 d**) with the instrumentation detailed in **Methods**. We induced acoustoelectric difference frequencies within the range reported for motor stimulation^22,25^, by applying an electric field that was close in frequency to the propagating acoustic field. We found that acoustoelectric amplitudes increased as Δ*f* decreased (**Fig. 2 e**).

Then, a recovery surgery was performed to insert two platinum-iridium electrodes connected to a small plug at the back of the mouse head so that the signal in the brain at the motor cortex could be measured in 3 wild type mice (see **Methods** for surgery details), using the same platinum-iridium electrodes and spacing as the saline petri-dish experiments. After surgery recovery, each mouse was anaesthetized with ketamine/xylazine and the same electrode and applied fields as the petri-dish were applied to the mouse brain. We show representative acoustoelectric time-series traces for Δf = 1Hz and Δf = 4Hz (**Fig. 2 f i**), where the electrical difference frequency is present alongside a DC offset. The corresponding spectral peaks (**Fig. 2 f ii**) show the relative amplitude at 1Hz was higher than at 4Hz, when the applied electric and acoustic field amplitudes were the same. *In vivo* acoustoelectric amplitudes decreased with increasing Δ*f* (**Fig. 2 f iii**, n=3 mice), suggesting acoustoelectric neuromodulation would be most likely where the difference frequency between the applied fields was smallest.

### *In vivo* acoustoelectric DC neuromodulation

The same recovery surgery as described in **Fig. 2** was performed in 9 wild type mice, where each mouse was anaesthetized with ketamine/xylazine and the stimulation experiment conducted during the lightening anaesthesia window so that electromyography (EMG) responses could be measured in parallel to the evoked acoustoelectric field in the brain (see **Methods** for experiment details). Two platinum-iridium ring electrodes (diameter 2cm) applied the bipolar electric field, and the ultrasound transducer was positioned via stereotaxic instrument over the head of the mouse (**Fig. 3 a**) with two electrodes placed in the forepaw to measure neural evoked muscle responses (EMG), with electrically-attenuating acoustically-semi-transparent material (F21 characterized in **Supplemental Note 2**) placed between the transducer cone and gel creating an acoustic connection while reducing the electrical interference from the electrical transducer artefact^26^.

**Figure 3.**
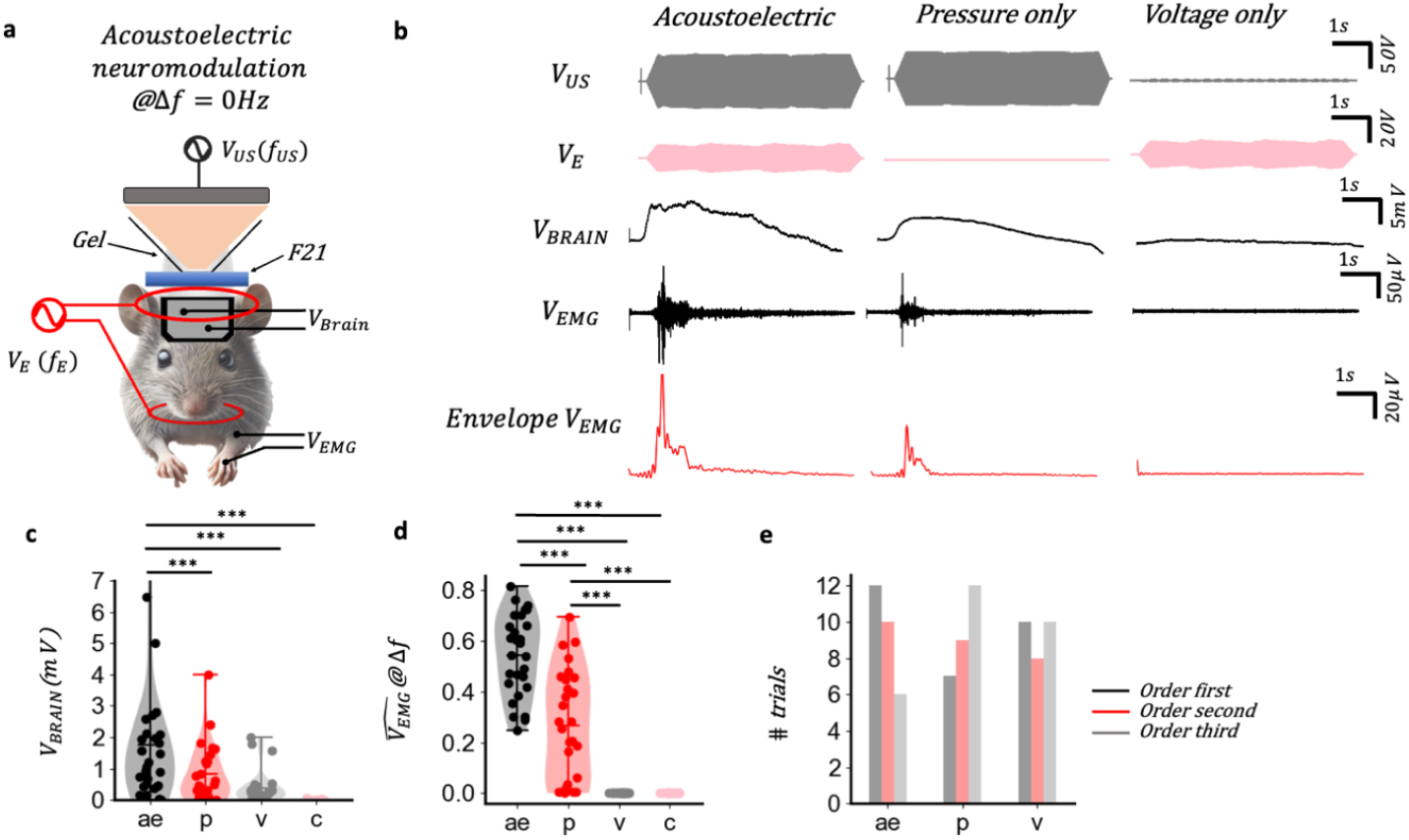
*In vivo* acoustoelectric amplitude neuromodulation @ Δ*f* = 0*Hz*. **a**, *In vivo* arrangement to apply ultrasound while electrically shielding the RF artefact from the ultrasound with F21, with applied bipolar electrical stimulus via external ring electrodes. Acoustic and electric field are applied at 500kHz. **b**, Example triplet with applied pressure 2.5MPa, voltage output 20V, showing *V*_*BRAIN*_, *V*_*EMG*_ and the envelope of *V*_*EMG*_. Acoustoelectric *V*_*BRAIN*_= 9.10*mVpp*, pressure only *V*_*BRAIN*_= 4.99 *mV* and voltage only *V*_*BRAIN*_ = 1.11*mV. V*_*EMG*_ software filtered between 100-1000Hz, Envelope of *V*_*EMG*_ low-pass filtered at 10Hz. **c**, triplets applied in randomized order to account for anaesthesia depth changes. *V*_*BRAIN*_ comparison between acoustoelectric, pressure only, voltage only and a no stimulation control. ANOVA F(3,27) = 8.50, p = 4.8e-5, post hoc comparisons using the Tukey HSD test, (AE–P, p = 0.02; AE-V, p = 0.0003; P-V, p = 0.52; AE-C, p = 0.00; P-C, p = 0.21; V-C, p = 0.80); over 9 mice, 28 triplets in total. **d**, Same data set as in c) comparison of Hilbert EMG height normalized across the triplet denoted by 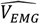. Statistics on the log ratios; ANOVA F(3,27) =252.16, p = 0, post hoc comparisons using the Tukey HSD test, (AE–P, p = 0.002; AE-V, p = 0.0; AE-C, p = 0.0, P-V, p = 0.0, P-C, p = 0.0, V-C, p = 1.0). **e**, Order analysis showing number of acoustoelectric (AE), pressure (P) and voltage only trials and their experimental order. Chi-squared test X^2^ (4, *N* = 28) = 5.5, *p* = .23; Bonferroni corrected pairs; AE–P, p = 0.19; AE-V, p = 0.45; P-V, p = 0.54.

A series of three tests were performed to determine the effect of applying an acoustic or electric field alone and then both together, referred to as a triplet. Within each triplet the voltage and pressure were the same, and each trial was 6 s long with a 0.25 s onset and offset ramp. The electric potential induced in the brain at Δ***f*** estimates the strength of the neuromodulatory electrical field, and the EMG potential in the paw estimates the strength of the evoked motor response. When both fields were applied together and at the same frequency (500kHz), an acoustoelectric difference frequency Δ***f*** was measurable in the brain at 0Hz (DC). This low frequency electric field typically had a steep gradient at the onset shown by the representative recordings in **Fig. 3 b** and **Video Supplement 1**. Simultaneous EMG recording shows the direct muscle response at the onset and how it differs between the three different stimuli, with the EMG envelope computed via Hilbert transform and then low pass filtered at 10Hz.

To overcome the changing sensitivity to the applied stimulus due to the anaesthesia depth changes, we applied each triplet in counterbalanced order (see **Supplemental Note 3** on impact of anaesthesia on evoked EMG amplitudes) over 9 mice with 3 or 4 triplets per mouse yielding a total of 28 trials, where the only data selection criterion was that the anaesthesia should be light enough to evoke an EMG response in any trial of the triplet. An important additional question is whether the acoustic or electric fields induced a neuromodulation response alone. We found that the DC electric potential induced in the brain by the acoustoelectric stimulation was larger than that induced by the acoustic or electric stimulation alone, and larger than a no stimulus control (ANOVA F(3,27) = 8.50, p = 4.8e-5; **Fig. 3 c**). Similarly, the normalized EMG amplitude^27^ induced in the paw by the acoustoelectric stimulation was larger than the those induced by acoustic or electrical stimulation alone (ANOVA F(3,27) =252.16, p = 0; **Fig. 3 d**). The EMG response was not dependent on the stimulation order (Chi-squared test X^2^ (4, *N* = 28) = 5.5, *p* = .23; Bonferroni corrected pairs; AE–P, p = 0.19; AE-V, p = 0.45; P-V, p = 0.54; **Fig. 3 e**).

These results suggest that applying an acoustic and electric field at the same frequency acoustoelectrically induces a DC electric field that is sufficiently strong to modulate neural activity.

### *In vivo* acoustoelectric AC neuromodulation

We next tested the neural response to acoustic and electric fields applied at a difference frequency Δ*f* of 1 Hz, i.e *f*_*US*_=500 kHz, *f*_*E*_=500.001 kHz (n=9 mice, 3-6 trials per mouse, total of 45 trials), where trials that occurred before any EMG response was possible were excluded, and the trial number variability was due to the experiments ending at different times based on when spontaneous mouse movements occurred. The difference frequency (Δ*f* =1Hz) during acoustoelectric stimulation was present in both the signal recorded in the brain and the envelope EMG, but was not present when the acoustic or electric field was applied alone as shown in the representative recording **Fig. 4 a**.

**Figure 4.**
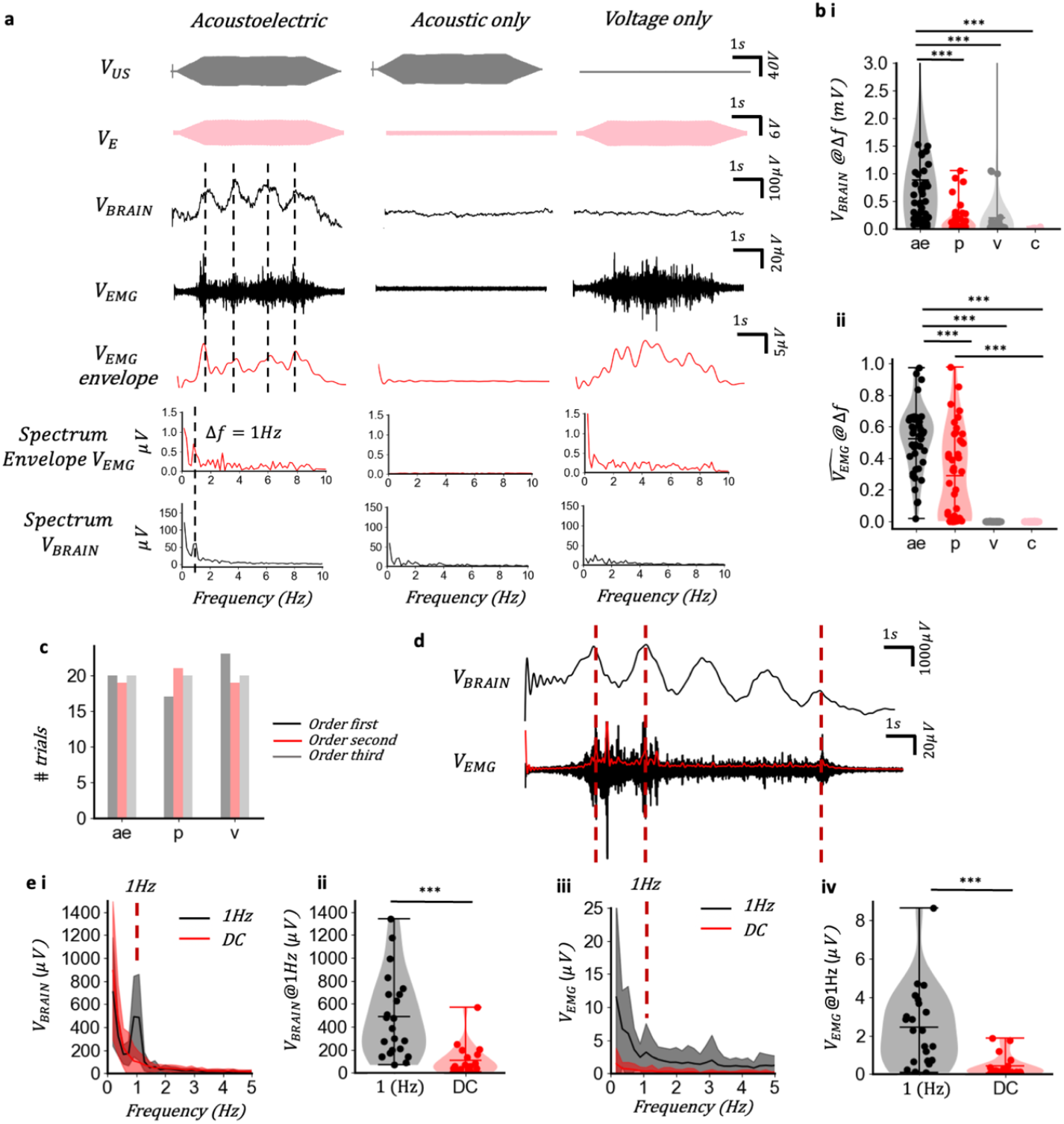
*In vivo* acoustoelectric neuromodulation @ Δ*f* = 1*Hz*. **a**, triplet where the acoustic wave was applied at 500kHz 1.8MPa, and the bipolar voltage output stimulus at 12Vpp, 500.001kHz. Δ*f* = 1*Hz* at *V*_*BRAIN*_ is mirrored in the envelope of *V*_*EMG*_ and reflected in the spectrum. Using the spectrum to measure amplitude, acoustoelectric = 510 *μV*, pressure = 80 *μV*, and voltage = 60*μV*. **b**, i) The spectrum amplitude at 1Hz was calculated for 9 mice, with a total of 45 triplet tests performed when the anaesthesia was light enough to evoke an EMG response. Spectrum was computed on the central 4 seconds to exclude the onset and offset ramps. ANOVA F(3,44) = 6.36, p = 0.0004, post-hoc comparisons using Tukey’s HSD test, (AE–P, p = 0.002; AE-V, p = 0.004; P-V, p = 0.5; AE-C, p = 0.02; P-C, p = 0.9; V-C, p=0.9); ii) The spectrum *V*_*EMG*_ amplitude at 1Hz ratio per triplet was compared using the log of the ratio for statistical tests. ANOVA F(3,43) = 370.69, p = 0, post-hoc comparisons using the Tukey HSD test, (AE–P, p = 2.27e-5; AE-V, p = 0.00; P-V, p = 0.00; AE-C, p = 0.00; P-C, p = 0.00; V-C, p=1.00); **c**, Experimental order analysis. **d**, Time series *V*_*BRAIN*_ were time correlated to the envelope *V*_*EMG*_ peaks(red). Shown are 3 time-correlated peaks demonstrating both the correlation and irregularity of the *V*_*EMG*_ **e**, Frequency specificity comparison between *V*_*BRAIN*_ at 1Hz and 0Hz, where *V*_*BRAIN*_ has selection criteria such that the 1Hz amplitude is larger than the 0.5Hz amplitude eliminating trials where electrochemical or pressure onset effects dominated. i) *V*_*BRAIN*_ comparison at Δ*f* = 0 and 1Hz, with mean and s.d. error bar. ii) T-test comparison between Δ*f* = 0,1Hz acoustoelectric *V*_*BRAIN*_ measurements, two sided t_(21)_= 4.38, p = 8.5e-5; *V*_*BRAIN*_at 1Hz (mean±s.d. = 491±351 µV); *V*_*BRAIN*_at 0Hz (mean±s.d. = 108.2±130µV). iii) *V*_*EMG*_ comparison between 0, 1Hz datasets shows 1Hz peak only in Δ*f* = 1*Hz* data set iv) T-test comparison between Δ*f* = 0,1*Hz* acoustoelectric *V*_*EMG*_, two sided t_(21)_= 4.11, p = 0.00019; 1Hz *V*_*EMG*_, (mean±s.d. = 2.45±2 µV); 0Hz *V*_*EMG*_, (mean±s.d. = 0.42±0.55µV).

Using the same counterbalanced experimental protocol as the acoustoelectric DC amplitude experiments, triplets were performed over the same 9 mice, with 4-5 repeats per mouse yielding a total of 44 triplets. To quantify the amplitude of the acoustoelectric signal at Δ*f* = 1*Hz*, we computed the spectrum using Fourier transformation and extracted the amplitude at the 1 Hz bin. We found that the 1Hz electric potential induced in the brain by the acoustoelectric stimulation was larger than those induced by the acoustic or electrical stimulation, or a no stimulation control (ANOVA F(3,44) = 6.36, p = 0.0004; **Fig. 4 b i**). Similarly, the normalized EMG amplitude induced in the paw by the acoustoelectric stimulation was larger than those induced by the acoustic or electrical stimulation (ANOVA F(3,43) = 370.69, p = 0; **Fig. 4 b ii**). The EMG response was not dependent on the stimulation order (Chi-squared test X^2^ (2, *N* = 44) = 2.84, *p* = 0.24; Bonferroni corrected pairs AE–P, P = 0.83; AE-V, P = 0.27; P-V, P = 0.36; **Fig. 4 c**).

The EMG response at 1 Hz was often irregularly shaped, with significant DC onsets potentially due to; acoustic standing waves in the mouse head (see **Supplemental Note 4**), electrochemical offsets at the electrode interface (**Supplemental Note 5**), and the onset response induced by the acoustic field alone (see **Fig. 3 b**, acoustic only signal). In a typical representative signal (**Fig. 4 d**) we found two time-correlated peaks at the beginning and one at the end. Shown in the **Video Supplement 2**, the variance in the acoustoelectrically generated amplitude in the measured brain signal *V*_*BRAIN*_ can be seen, with the mouse responding via paw movements to only the largest voltage changes in the brain.

To validate that the 1Hz spectral amplitude was not confounded by the DC offsets observed in some recordings, we repeated the analysis but with only the acoustoelectric recordings where the DC offset did not dominate the 1Hz acoustoelectric signal. We found an evident peak at the 1 Hz difference frequency in both the electric signal recorded in the brain (**Fig. 4 e i-ii**) and the paw EMG (**Fig. 4 e iii-iv**) that did not exist when the US and electric fields were applied at the same frequency, i.e. DC acoustoelectric stimulation (amplitude spectral density (ASD) at 1Hz vs 0Hz: *V*_*BRAIN*_: t(21)= 4.38, P = 8.5e-5; *V*_*EMG*_: t(21)= 4.11, P = 0.00019). Therefore, the acoustoelectric field generated in the brain affects the frequency of evoked neuromodulation.

### Acoustoelectric artefact tests

To determine if temperature dissipation might be the cause of the response during acoustoelectric neuromodulation, we applied an acoustoelectric 0.5Hz difference frequency where the acoustic field was 500kHz and the applied voltage 500.0005kHz (**Fig. 5 a i, ii**). A temperature probe was placed in the ultrasound gel above the skull and below the end of the ultrasound cone while a continuous 6 second acoustoelectric signal was applied, repeating 6 times for pressure or electric field alone, and electric and pressure field together to generate an acoustoelectric difference frequency at 0.5Hz. While applying continuous ultrasound at 1MPa does increase the temperature (**Fig. 5 b**, shown in blue), the acoustoelectric and pressure only trials had a very high mean temperature correlation (0.98; Pearson, **Fig. 5 b**), and no significant group difference (AE-P = 0.9, n=36; **Fig. 5 c**). This suggests the pressure and acoustoelectric trials are not differentiated by temperature, unlike the EMG and brain signal differences in the acoustoelectric versus pressure only experiments shown in **Fig. 4**. Hence, temperature is unlikely to be the cause of acoustoelectric neuromodulation.

**Figure 5.**
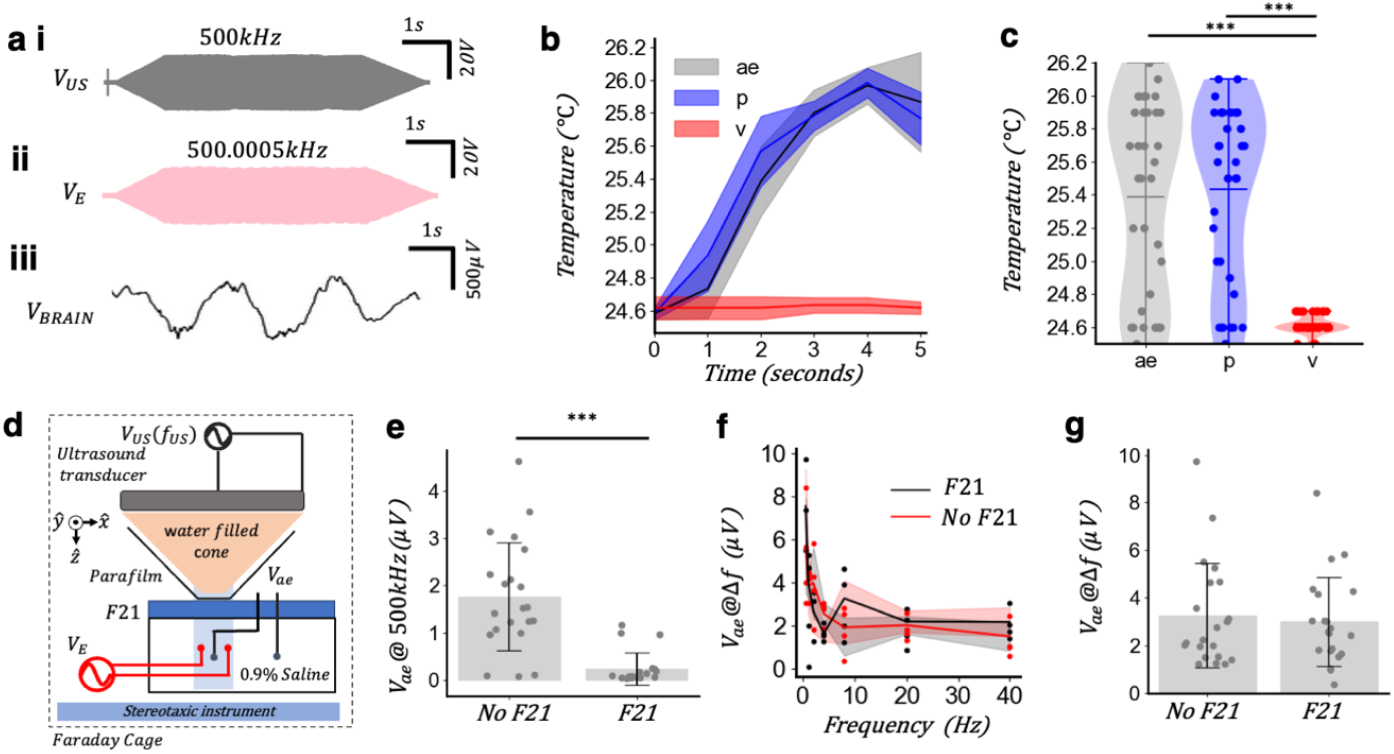
Acoustoelectric artefact tests. **a**, 1MPa pressure, 40V pp applied voltage, 6 repeats per AE/P/V group, using a thermometer placed in the ultrasound gel just above the mouse skull and below the ultrasound cone. Δ*f* = 0.5Hz. i) *V*_*US*_ at 500kHz, 1MPa signal applied in each recording shown ii) *V*_*E*_ at 20V. iii) *V*_*BRAIN*_ filtered <10Hz revealing 0.5Hz acoustoelectric oscillation. **b**, Temperature change as a function of time since beginning of trial with s.d. error shown for each AE/P/V group. **c**, ANOVA F(2,35) = 35.6, p = 1.5e-12, post-hoc comparisons using Tukey’s HSD test, (AE–P, p = 0.9; AE-V, p = 9e-10; P-V, p = 5.9e-11); **d**, Saline petri dish within the electrophysiology set up to induce and measure an acoustoelectric field. **e**, T-test comparison of the amplitude of the electric field generated by the ultrasound transducer at 500kHz, measured in the medium with and without F21, two sided; (t_(20)_= 5.72, p = 1.15e-6), note that since the electric field at the carrier was large, these carrier amplitudes are measured after 5kHz −3dB point low pass filtering on the preamplifier front end. **f**, Varying Δ*f* with and without F21 with 4 repeats at each frequency. **g**, T-test comparison of the Δ*f* acoustoelectric field amplitude with and without F21 using independently applied electric field, two sided; (t_(20)_= 0.38, p = 0.70).

To eliminate electric field only mixing between the transducer artefact electric field (*V*_*US*_) and the independently applied electric field (*V*_*E*_) as the cause of the difference frequency, we calibrated an acoustoelectric field in a saline petri-dish to have the focal maximum at the measurement electrodes (**Fig. 5 d**, with calibration procedure and spatial map described in **Supplemental Note 6**). The electric signal powering the ultrasound created an electric artefact at the pressure frequency which could be measured in the saline petri dish and attenuated by inserting electrically insulating F21 material (**Fig. 5 e;** p = 1.15e-6). An acoustoelectric field was measured with and without the electrically attenuating F21 material, at a series of electric field difference frequencies to induce a spectrum (**Fig. 5 f**). There was no difference in the acoustoelectric amplitudes between the electrically attenuating F21 and no F21 group (**Fig. 5 g;** p = 0.38), suggesting that the induced acoustoelectric signal did not originate from mixing with the electric field artefact from the ultrasound transducer.

Non-linearity in recording hardware could also generate an electric difference frequency. To test it, we fed driving voltage signals of the electric and acoustic stimulation to the input of the recording preamplifier via an isolation transformer. In this case, there were no recordings at the difference frequency, suggesting the acoustoelectric signal did not artefactually originate from frequency mixing in the recording hardware (**Supplemental Note 7**).

### *In vivo* transcranial ultrasound stimulation is impacted by the electrical artefact powering the transducer

Now that we have evidence that acoustoelectric neuromodulation is possible when an independent electric field and a propagating acoustic field are applied to an ionic medium, we investigated the impact of the electric field artefact generated by powering the ultrasound transducer, to determine if it may also interact with the propagating acoustic field in ultrasound stimulation.

To determine the origin of the electric signal present in the mouse brain at the acoustic frequency, we first placed an acoustically transparent and electrically insulating material (2mm thick F21, Precision Acoustics, UK; characterized in **Supplemental Note 2**) between the end of the ultrasound cone and the mouse. We found the electric signal at the acoustic frequency was attenuated (**Supplemental Note 8 a-c**). We then electrically isolated the transducer from ground by inserting a switch into the cable connecting the transducer to the RF amplifier, which massively increased the amplitude of the electric signal transmitted into the mouse (**Supplemental Note 8 d-f**) as the energy injected into the transducer needed to find a path to ground via nearby ionic media such as the mouse. These results suggest the electrical artefact from powering the transducer capacitively transmits electrical energy into nearby ionic media, similar to an electrical antenna^27^. Using a saline phantom, we then verified that electrical sum and difference frequencies emerge with the focality of the ultrasound when a pulsed pressure signal is applied alone (**Supplemental Note 9 a-b**). The amplitude of these focal frequency mixing products was attenuated when electrically shielding, acoustically transparent F21 was placed between the transducer cone and the saline (**Supplemental Note 9 c-d**).

To decouple the ultrasound acoustic field and electric artefact *in vivo*, we performed an F21 electrical shielding experiment (**Fig. 6 a**) with a pulsed ultrasound stimulation protocol where the pulse was on 0.5s and off for 1.5s (25% duty cycle) at 500kHz as these parameters have been reported to be highly effective at eliciting an evoked ultrasound stimulation responses previously^28^. During the lightening anaesthesia window 2mm thick F21 was inserted into the gel below the ultrasound cone to shield the electrical artefact coming from the ultrasound transducer, decreasing the amplitude of the electric field present in the medium, and then removed and repeated 1-2 times per mouse until spontaneous movements started, alternating the experimental order to minimize anaesthesia depth confounds (see **Supplemental Note 3**). To overcome the small acoustic attenuation induced from the F21 (**Supplemental Note 2**), 15% was added to the pressure output when the F21 was in place in all experiments.

**Figure 6.**
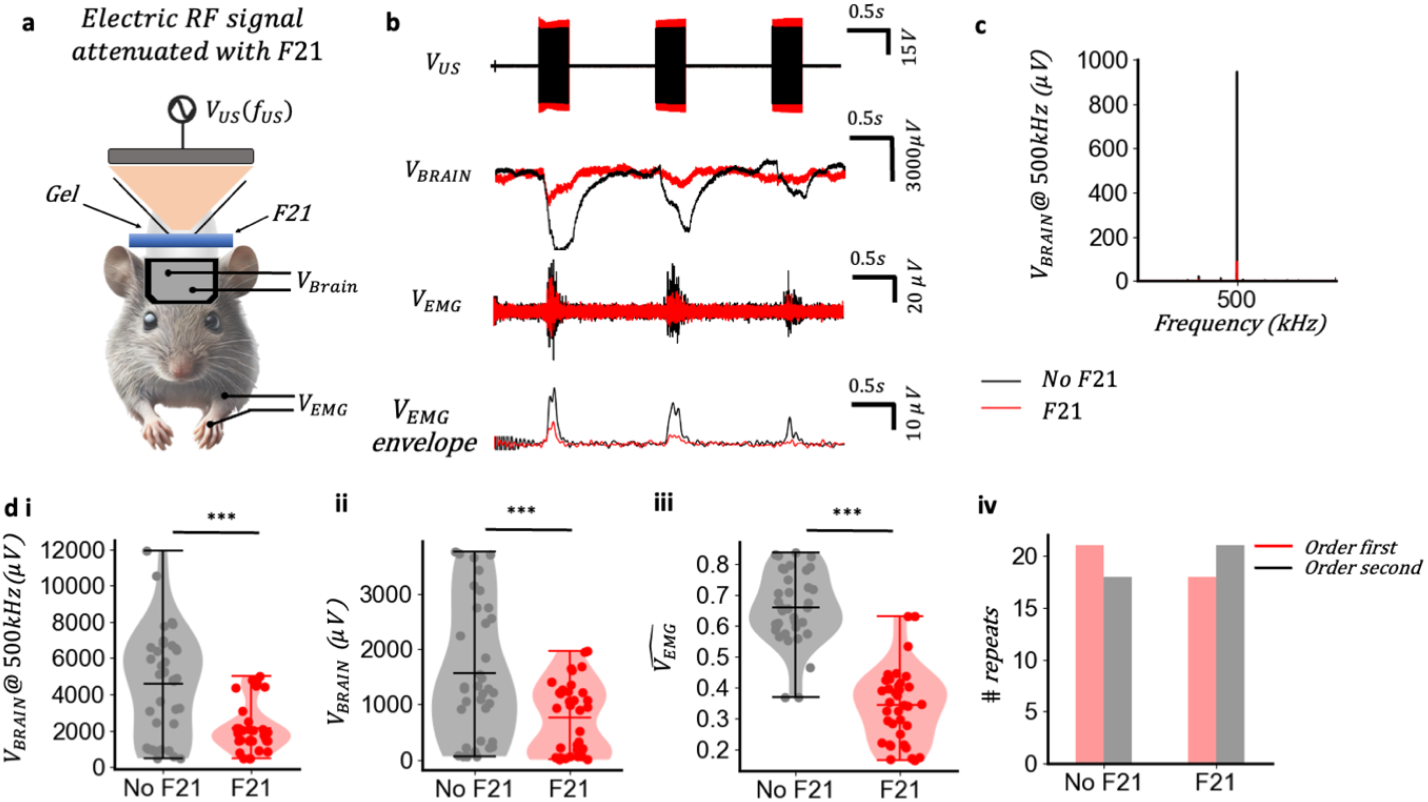
*In vivo* ultrasound brain stimulation with attenuation of the electrical artefact from the ultrasound transducer. **a**, Mouse undergoing pulsed ultrasound stimulation with F21 electrical attenuation material in place. Mice performed the with and without F21 test twice. **b**, Representative with (red) and without (black) F21 experiment, with *V*_*US*_ shown 1kHz low pass filtered at *V*_*BRAIN*_, *V*_*EMG*_ and envelope *V*_*EMG*_. Pressure without F21 = 2MPa; pressure with F21 = 2.4MPa. **c**, Representative amplitude spectrum at *V*_*BRAIN*_ comparison between electric artefact with and without F21. **d**, i) carrier amplitude comparison; t-test, two sided *t*_(01)_= 4.65, p = 1.47e-5; F21 group (mean±s.d. = 2895 ± 1253*μV*); without F21 group (mean±s.d. = 4601 ± 2116 *μV*) n=7mice, 36 trials, during light ketamine/xylazine anaesthesia. ii) *V*_*BRAIN*_ comparison log normalized to take pair based pressure differences into account; t-test, two sided *t*_(01)_= 3.52, p = 7e-4; F21 group (mean±s.d. = 755 ± 630*μV*) No F21 group (mean±s.d. = 1572 ± 1217 *μV*) using data as in d. iii) *V*_*EMG*_ normalized per pair to account for changing anaesthesia; t-test, two sided on log normalized pair; *t*_(01)_= 4.65, p = 1.4e-5; F21 group (mean±s.d. = 0. 34 ± 0.11*μV*); No F21 group (mean±s.d. = 0. 665 ± 0.11 *μV*) using data as in d. iv) Experimental order analysis.

In **Fig. 6 b** we show a typical representative example, where we applied a pulsed ultrasound signal without F21 and saw it produced a larger amplitude low frequency brain signal than the group with F21 in place at 2.4MPa. The resulting EMG signal measured without F21 has larger spikes, while electrically shielding the transducer with F21 yielded smaller EMG responses (shown in red in **Fig. 6 b**, shown with simultaneous video and measurements in **Video Supplement 3**). EMG responses also reduced on the third pulse as compared to the first pulse, and the low frequency electrical signal measured in the brain (*V*_*BRAIN*_) was not always the same amplitude per pulse, despite the pressure output being the same amplitude for each pulse in a trial. Looking at the spectral amplitude of the electrical artefact from the ultrasound in the same representative sample (**Fig. 6 c**), shows F21 successfully attenuated this signal in the mouse brain. At the group level, with 7 mice yielding a total of 36 pulses with F21/noF21 pairings, the electrical insulation properties of F21 attenuate the electrical carrier artefact from the transducer (**Fig. 6 d i**, p = 1.47e-5). This trend is mirrored in the pair normalized brain signal amplitude comparison (**Fig. 6 d ii**, p = 7e-4), as well as the EMG signal amplitudes (**Fig. 6 d iii**, p = 1.4e-5).

Finally, to ensure that the order of trials was not responsible for the increased amplitude when no F21 was present we performed a Chi-squared test on the counterbalanced data showing there was no significant relationship between the order of trials and the amplitude of the EMG response X^2^ (3, *N* = 35) = 1.32, *p* = 0.24 (**Fig. 6 d iv**). Hence *in vivo* ultrasound stimulation responses can be attenuated by reducing the amplitude of the electric artefact in the medium, independent of time-based anaesthesia depth changes.

### *In vivo* ultrasound stimulation is not due to electric field only mixing

To determine if the stimulation effect may be due to the electric field artefact from the 500kHz electric field applied by the ultrasound transducer mixing with itself at the measurement electrodes, we performed an acoustic disconnection test where the electric field remained connected via gel connection with the ultrasound cone, but the acoustic field was removed by leaving an air gap below the end of the cone (**Fig. 8 a**). To overcome a slight reduction in the amplitude of the electric field when a smaller volume of conductive gel was present, we added 15% to the output pressure.

The representative plots (**Fig. 7 b ii**) show the voltage recorded at the measurement electrodes in the brain (*V*_*BRAIN*_), where no EMG was evoked in the electric artefact only case while the acoustic field did induce a response (**Fig. 7 b iii, iv**, and **Video Supplement 4**). This was repeated across 6 mice, with 2 trials of each case recorded for each mouse, each with 3 pulses recorded. The carrier frequency measured in the brain was similar between acoustically connected (no gap) and disconnected (gap) groups, though the larger surface area gel connection enabled better electrical transmission (**Fig. 7 c**; p=0.0018). There were no DC offsets in the brain when there was an air gap blocking the acoustic signal (**Fig. 7 d**, p=8.8e-6), and no EMG responses were evoked in any trial (**Fig. 7 e**, p=6.0e-18). The difference between the measurement with and without the air gap was not affected by the measurement order (**Fig. 7 f**).

**Figure 7.**
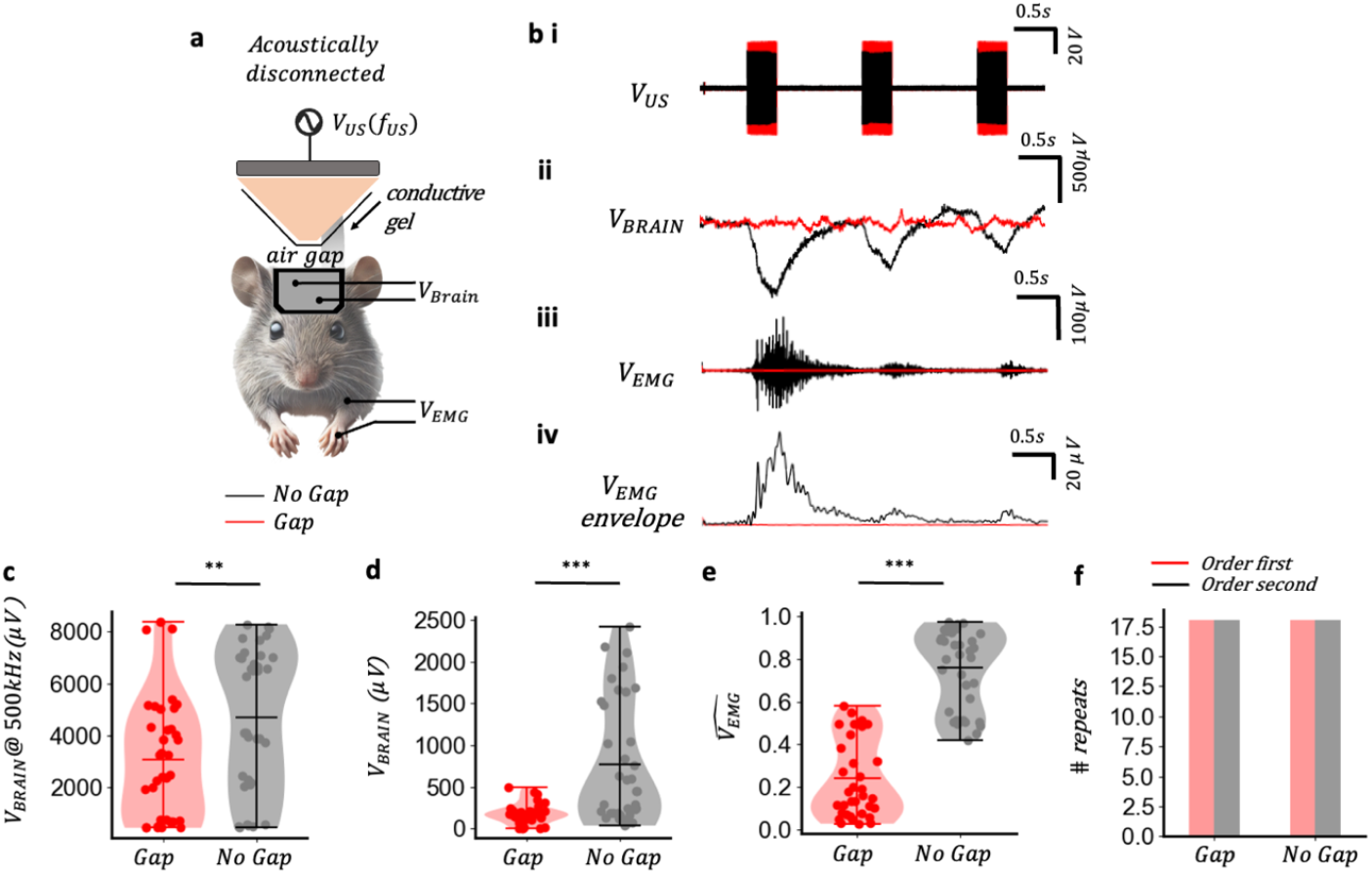
Ultrasound stimulation is not due to electric mixing. **a**, Instrumentation arrangement with acoustically disconnected measurement in mouse brain such that the acoustic wave is blocked by an air gap and the transducer remains electrically connected to the brain via conductive gel. 6 second recordings with 0.5second 500kHz 1MPa acoustic pulses. **b**, i) RF output signal for 1MPa pressure showing acoustically connected amplitude (black) 20% lower than acoustically disconnected amplitude. ii) low frequency brain signal *V*_*BRAIN*_, (<1kHz) comparing no gap/acoustically connected (black) against electrical connection only/air gap (red). iii) EMG trace *V*_*EMG*_ (100Hz-1kHz) for both no gap (black) and gap (red) cases. iv) Hilbert EMG traces (<10Hz) for no gap (black) and gap. **c**, Carrier amplitude as measured via the brain electrodes for gap (red) and no gap (black). *t*_(01)_= −3.23, p = 0.0018; gap (mean±s.d. = 3066±2302); no gap (mean±s.d. = 5967±4773) n=6 mice, with 2 trials each with 3 pulses for each mouse yielding total 36 pulses. **d**, Brain height comparison between gap (red) and no gap (black), *t*_(01)_= −4.79, p = 8.81e-6; gap (mean±s.d. = 174.2±117.2); no gap (mean±s.d. = 771.2±726.9) n=6 mice. **e**, Comparison between np gap and gap EMG across n=6 mice, with 2-3 trials per mouse, normalized across each trial to account for time based variation, statistics performed on log on normalized triplet; t_(01)_= −9.06, P = 2e-13; gap (mean±s.d. = 0. 24 ± 0.18); no gap (mean±s.d. = 0. 75 ± 0.18); n=6 mice. **f**, Order analysis X^2^ (3, *N* = 35) = 0.16, *p* = 0.68.

These results support the evidence that the DC electric field present in the brain tissue is generated by an acoustoelectric interaction between the ultrasound and its electric artefact.

## DISCUSSION

Non-invasive electrical brain stimulation shows promise as an alternative to invasive neurosurgical approaches for treating epilepsy^29^, obsessive compulsive disorder^30^ (OCD), depression^31^, Alzheimer’s disease^32^, and for inducing neurogenesis after^33^, while avoiding the risks of surgical intervention. However, clinical translation requires techniques that deliver better targeting, controllability and safety. Ultrasound stimulation offers high spatial specificity and depth, yet uncertainty about its mechanism persists^34,35^. A modality that harnesses the well-characterised electrical models of neural signalling while retaining the spatial focality of ultrasound would enable independent optimisation of amplitude and safety through acoustic and electric control axes.

Acoustoelectric neuromodulation achieves this via a non-linear interaction between an applied electric field and a focused ultrasound field, generating a focal difference-frequency electric field that can drive neuromodulation. The observed amplitude and frequency dependence of responses on the acoustoelectrically generated difference frequency argues against pressure-only mechanisms as the primary driver. For example, acoustic radiation force^36^ (ARF) can be dissociated from the difference-frequency response by varying the applied electric-field frequency: when a 1 Hz difference frequency is generated and evokes stimulation at 1 Hz, ARF alone cannot account for the effect because the response also requires the independently applied high-frequency electric field. Taken together, the results argue against pressure-only accounts - including non-linear acoustics^37^, acoustically induced thermal dissipation^9,38^, cavitation^39^, or auditory pathways^10,40^ - as sufficient explanations.

Acoustoelectric neuromodulation is also not explained by electric mixing only from the transducer drive and an independent applied electric field. The artefactual electric field produced by powering the transducer is not directly proportional to the acoustoelectric difference-frequency signal, based on testing with an electrically attenuating material. Pressure-induced vibration of recording electrodes, which could generate non-linear electrochemical artefacts ^41^, is likewise unlikely: free-field saline phantom experiments showed that acoustoelectric amplitude depends on the geometry of the applied fields in the medium^17^, inconsistent with a simple vibration artefact.

Because the ultrasound drive voltage can capacitively couple into surrounding ionic media, an artefactual electric field coexists with the propagating acoustic field during ultrasound stimulation. Since both occur at the same frequency, their interaction yields a difference frequency at 0 Hz (DC). The dependence of response amplitude on the electric-field magnitude present in the medium when acoustic and electric frequencies are matched (**Fig. 3**) indicates that acoustic-only mechanisms - non-linear acoustics^37^, thermal dissipation^9,38^, cavitation^39^, and auditory pathways^10,40^ - cannot solely account for ultrasound-evoked neuronal activation. Conversely, electric-field–only mixing produced by the transducer drive did not elicit stimulation, arguing against purely electrical or electrochemical origins. Together, these observations indicate that both acoustic and electric fields are required to generate a focal difference-frequency electric field and that the acoustoelectric interaction makes a substantial contribution to ultrasound brain stimulation.

Limitations include the relationship between mouse head size and acoustic wavelength, which promoted non-focal standing waves (**Supplemental Note 4**). Using higher-frequency transducers or larger heads (e.g., humans) should reduce standing-wave artefacts and yield more focal acoustoelectric fields^42^. We also exceeded the recommended thermal index for human use^43,44^; translating this approach will require waveform optimisation and careful consideration of skull attenuation and aberration^45^.

Acoustoelectric neuromodulation offers advantages over ultrasound alone. It leverages the extensive literature on how electric fields influence neurons^46^ and allows rapid *in vitro* optimisation of focused electrical parameters. With acoustic and electric inputs as independent control axes, safety and efficacy can be tuned separately.

Future work should further isolate and quantify the respective acoustic and electric contributions *in vivo*. Ultrasound responses depend on the electrical environment, including grounding, transducer geometry, and the electric field necessary to drive the transducer (e.g., focused transducers generally require lower drive voltages than planar transducers to achieve equivalent pressures ^47^); endogenous electric fields may also shape the net field inside neural tissue. More broadly, acoustoelectric interactions are likely to influence related modalities that combine acoustic and electromagnetic fields, including magneto-acoustic neuromodulation with static fields^48^, MR-guided tFUS^34^, and conventional ultrasound brain stimulation.

Our results demonstrate that acoustoelectric mixing can directly elicit neuromodulation and constitutes a key mechanism underlying ultrasound-induced brain stimulation. By clarifying how acoustic and electric fields interact within neural tissue, this work advances spatially precise, non-invasive control of brain activity.

## METHODS

### In-vivo mouse experiment

*Animals*. Mice were wild type C57BL/6 male and female mice, aged between 3-6 months. Mice were housed in standard cages in Imperial College London animal facility, with ad libitum food and water in a controlled light-dark cycle environment, with standard monitoring by veterinary staff. The Imperial College of London’s Animal Welfare and Ethical Review Body approved all animal procedures, and all experiments were performed in accordance with relevant regulations/according to the United Kingdom Animals (Scientific Procedures) Act 1986.

*Surgery*. Mice were anaesthetized using 1-3% (vol/vol) isoflurane in oxygen and subcutaneously administered Vetergesic (0.1mg/kg) and Carprofen (5mg/kg) to inhibit pain and inflammation and saline for hydration (10ml/kg/hour). A Neurotar compatible head plate was attached to the mouse using clear dental cement (PalaXpress Dental Cement, AgnTho’s AB, Sweden) and a craniotomy performed with a drill (diameter 0.5mm) at the intended electrophysiological measurement sites. The position of the craniotomy relative to bregma was anteroposterior 0.0 mm, mediolateral −2 mm for the motor cortex electrode and anteroposterior −3.5mm, mediolateral 2.25mm for the visual cortex reference electrode used to measure electrical amplitudes within the motor cortex. The recording electrodes were 98% platinum-iridium wire segments (0.5mm diameter wire, VWR, Lutterworth, UK) that were positioned over the drill holes so that the end of the wire was touching the brain, and dental cement secured it to the head bar. Nail polish was then applied to the electrodes to electrically insulate the platinum-iridium wire, so that only the portion of wire located in the brain was exposed. Low toxicity silicone sealant (Kwik-Cast, World Precision Instruments) was then poured around the electrodes to seal moisture around the craniotomy site and over the exposed skull, whilst also stabilizing the electrodes in position. The mouse was then recovered in a warming chamber and administered Carprofen (5mg/kg) for three days after surgery with daily weight checks, then allowed to recover for one week before experiments commenced.

#### Ultrasound fields application

The acoustic field was applied using a curved ceramic (PZT) 500 kHz ultrasound transducer (60mm diameter, 63.5 mm acoustic path length; Precision Acoustics Ltd, UK). The ultrasound transducer was driven by an arbitrary function generator (Handyscope HS5, TiePie engineering, Netherlands) and a 40W linear power amplifier (240 L, Electronics & Innovation Ltd). The ultrasound transducer was mounted in a cone and filled with water characterized in **Supplemental Note 1**. Non-conductive Parafilm was stretched over the end of the cone to seal the water in, while the transducer was positioned via stereotaxic instrument above the mouse head. F21 was inserted in the gel to attenuate the electrical signal in the mouse brain emanating from the ultrasound transducer. Ultrasound gel (Anagel, UK) was applied on top of the mouse head, surrounding all electrodes, and providing an acoustic connection between the end of the ultrasound cone and the top of the mouse skull.

#### Electric fields application

The electric field was applied using a pair of 98% platinum-iridium wire electrodes (Alfa Aesar, 0.25mm diameter), that were formed into loops to increase the exposed surface area, further decreasing electrochemical induced offsets when the bipolar electric field was applied. One loop was gently inserted into the mouth (loop diameter 1cm), and the second located above the head (loop diameter 2cm), placed between the ultrasound cone and mouse head within the acoustic gel interface. The electrodes were driven by an arbitrary function generator (WiFiScope WS5, TiePie engineering, Netherlands) and a custom-made high-frequency voltage source made of glass core transformers (Hitachi Metals) with a maximum ±20 V amplitude at an effective 3 dB bandwidth range of 5 kHz to 2 MHz. To output voltages which surpassed the 12V function generator range, a 20W linear power amplifier was used (325LA, Electronics & Innovation Ltd). The transformer’s isolation prevented extraneous tissue charging and minimised total harmonic distortion.

#### Electrophysiology recordings

The electrode implant was connected to an SR560 preamplifier (Stanford Research Systems, USA) via a 1.27mm Dupont header which was cemented onto the back of the head bar. A low pass filter (5kHz −3dB frequency with circuit diagram in **Fig. S12**) was added whenever a high-frequency electric field was applied to prevent overloading the preamplifier.

#### Electromyography (EMG) recordings

EMG was measured using stainless steel EMG electrodes (0.3mm 622 diameter, Arbitter, UK) that were placed in the front paw, with the second electrode at the muscle innervation point on the forearm. The EMG electrodes were then connected to a differential amplifier (SR560, Stanford Research Systems) with a band pass filter (100-1kHz) to isolate just the EMG signal.

#### Data acquisition

All applied and monitored signals were recorded at 5MS/s, and were time synced using a combined multiple instrument (CMI) interface between the function generators and oscilloscopes with trigger inputs. The applied voltages, monitor channels and electric potentials were logged using data acquisition hardware (WiFiScope WS5, WiFiScope WS6 DIFF and Handyscope HS5, TiePie engineering, Netherlands). The logged signals were streamed from the data acquisition hardware to a workstation PC using custom C code which utilized the Tiepie SDK to control the triggers, channels, and function generator output. The compiled C code which called the oscilloscope commands was controlled through a Python wrapper script. An instrumentation diagram and connection details available in **Supplemental Note 11**. The low-pass filter described in **Supplemental Note 12**, was used in only the measurements with a large amplitude independently applied electric field (**Fig. 5** and below).

#### Experimental procedure

Anaesthesia of mice was achieved via subcutaneous injection of 75 mg/kg Ketamine and 10 mg/kg of Xylazine. Eye lubricant (Vaseline) was placed on the eyes to prevent them from drying out. The anaesthetized mouse was placed onto a thermal mat (Thermostar, RWD, China) to maintain homeostasis at 37°C and head position stabilized with the installed head clamp (Neurotar standard clamp, Finland) enabling precise stereotaxic localization of the ultrasound transducer over the head. The intracranial recording electrodes were connected to the recording amplifier, and the EMG electrodes were placed in the front paw and forearm and connected to the recording amplifier. Ultrasound gel (Anagel, UK) was applied on top of the mouse head, surrounding all electrodes to provide an acoustic connection between the end of the ultrasound cone and the top of the mouse skull. The stimulating loop electrodes were placed above the head and in the mouth as described above. The US transducer was positioned above the mouse head using a stereotaxic instrument. A F21 material was inserted into the gel to attenuate the US electrical artefact.

Acoustic and electric fields were then applied according to the instrumentation description (see **Supplemental Note 11 and 12**). Each experimental block included the three conditions (i.e., E, US, E+US), applied for 6 seconds with a 0.25-second ramp up and down periods in a counterbalanced order.

Measurements of EMG could only be made when the anaesthesia depth lightened approximately 45 minutes to an hour after administration. When small spontaneous movements started to occur, the experiment was ended, and the mouse was given Antisedan (1 mg/kg) subcutaneously to hasten Xylazine recovery and moved to a warming chamber for recovery. After recovering in the warming chamber under observation, the mouse was given water saturated food pellets and returned to their home cage.

Pressure amplitude ranged between 0.5-2.5MPa. Typical *in vivo* rodent experiments performed to date were done when the mouse is fully awake^49–52^, requiring lower pressure thresholds to evoke EMG and often with spontaneous movements also present. Here, experimental measurements were performed when the anaesthesia was light, and there were no spontaneous movements which required pressure magnitudes that were slightly higher than those measured in awake mice.

### Phantom measurements

*Petri dish phantom*. A small plastic petri dish (4cm diameter and 2cm depth) filled with 0.9% saline located where the mouse would be positioned within the Faraday cage used for *in vivo* electrophysiology experiments.

#### Tank phantom

A custom-built acrylic tank (50 cm× 20 cm × 20 cm) filled with 0.9% saline. An acoustic damping material (Aptflex-48) was attached to the back of the tank to minimise acoustic reflections. The phantom tank was covered with a Faraday cage.

#### US fields application

Similar to the *in vivo* experiment.

#### Electric fields application

Similar to the *in vivo* electric field application, but with the wire electrode forming short pins placed with a 10mm spacing around one of the recording electrodes.

#### Electrical recording

Similar to the *in vivo* intracranial electric field recording, with the electrode implant used for measurement and the same spacing as the *in vivo* experiments.

#### Pressure recordings

The acoustic field was measured in the tank phantom using a 0.2 mm needle hydrophone with a calibration of 52mV/kPa at 500 kHz provided by the manufacturer, Precision Acoustics Ltd and a DC-coupled preamplifier (Precision Acoustics Ltd).

#### Data acquisition

Similar to the *in vivo* data acquisition.

#### Procedure to identify US focus using the AE effect

To locate the acoustic focus in a saline phantom, an electric field was applied at 8kHz and the ultrasound transducer moved in 0.5mm increments to find the peak acoustoelectric amplitude at 492kHz. The acoustoelectric mixing amplitudes reached a maximum when the ultrasound focus was positioned over the voltage stimulation source (see **Fig. S6**). In the saline phantom, detailed calibration maps could be made like the ones shown (**Fig. S6 b i**,**ii**).

#### Spatial measurement procedure

For the 2-dimensional acoustoelectric free field maps recorded in a biomimicking tank phantom (20cmx20cmx50cms) filled with 0.9% saline, a recording was made for each pixel consisting of a ramped electric field at the beginning and end to avoid voltage spike transients in the preamplifier. To time sync individual recordings to one another, a half-amplitude 2 wavelength duration acoustic field was applied at the beginning of the signal. Once all measurements were made, each file was time aligned to the embedded acoustic field marker, after filtering to isolate the sum and difference frequencies, the individual files were extracted into a large three-dimensional array including time and two dimensions. This enabled the recovery of *ϕ*_*ae*_(*t*), such that we can obtain the time evolution of the generated 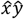 and 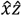 views.

### Signal processing and data analysis

Signal processing and data analysis were performed using custom Python scripts that utilised functions from the NumPy, SciPy, and Pandas Python libraries.

#### Data exclusion criterion

The only criterion for exclusion was when the anaesthesia was too deep to evoke any EMG response, despite the Δ*f* frequency being present in the brain (see **Supplemental Note 3**). All files were included in the dataset, even if there was no difference frequency sinusoid present due to poor data quality due to electrochemical offsets (**Supplemental Note 5**).

#### Computation of EMG envelope

EMG envelope was computed by performing a Hilbert transform and extracting the amplitude of the resulting analytical signal. A low-pass filter with a cutoff frequency of 10 Hz was applied to the resulting envelope.

#### Frequency domain analysis

A 1-D discrete Fourier Transform with Flat Top window to optimize amplitude accuracy was computed on the central 4 seconds of each recording, as we report amplitude spectral density (ASD) instead of power spectral density (PSD) to easily read the average amplitude of each measurement. After the Fourier transform is computed, the two-sided amplitude spectrum is multiplied by 2, and half the array is taken, converting it into its single-sided form.

#### EMG amplitude normalisation

The EMG amplitude (i.e., ASD at Δ*f*) at each stimulation condition was normalised to the total amplitude across the conditions at each triplet block to account for inter-block variation in responsiveness due to, for example, changes in anaesthetic level. Statistical tests were performed on the log values.

#### Peak alignment analysis

Peaks in the brain electric potential signal and EMG envelope signal were identified using the Python-based find_peaks function. An alignment between the peaks was defined as occurring when the EMG peak coincided with a peak of the brain signal, provided the time difference was within 1/5 of the period (i.e., 0.2 seconds for 1Hz stimulation).

### REPORTING SUMMARY

Further information on research design is available in the Nature Portfolio Reporting Summary linked to this article.

## DATA AVAILABILITY

The data supporting the results in this study are available within the paper and its supplementary information. The raw and analysed datasets generated during the study will be made publicly available before publication of the manuscript.

## CODE AVAILABILITY

Further information is available from the corresponding author upon request. Code supporting the findings in this article will be made publicly available before publication of the manuscript.

## AUTHOR CONTRIBUTIONS

J.L.R. conceived and developed the hardware, simulations and experiments and wrote and revised the paper. C.B. revised the paper. R.C. edited and revised the paper. N.G. edited and revised the paper and supervised experiments.

## ACKNOWLEDGEMENTS

N.G. was supported by the UK Dementia Research Institute (UK DRI)—an initiative funded by the Medical Research Council, Engineering and Physical Sciences Research Council (EPSRC) UK, Science & PINS Award for Neuromodulation, the NIHR IBRC Confident in Concept Award, and the American Alzheimer’s Association.

We thank all the members of CBS, Hammersmith at Imperial College London for their tireless care of the mice who live there, Dr. David Bono and Dr. Jon Howard for assisting with experimental set up, Dr. Patrycja Dzialecka for advice on mouse recovery surgeries, Dr. Hazael Montanaro for advice on acoustic finite element modelling simulation using Sim4Life, and Dr. Nawal Zabouri for initial rodent surgery training.

