## Supplementary Material for "Non-invasive in vivo acoustoelectric neuromodulation and its contribution to ultrasound stimulation"

### SUPPLEMENTARY MATERIALS

#### 1. Acoustoelectric and acoustic field characterization

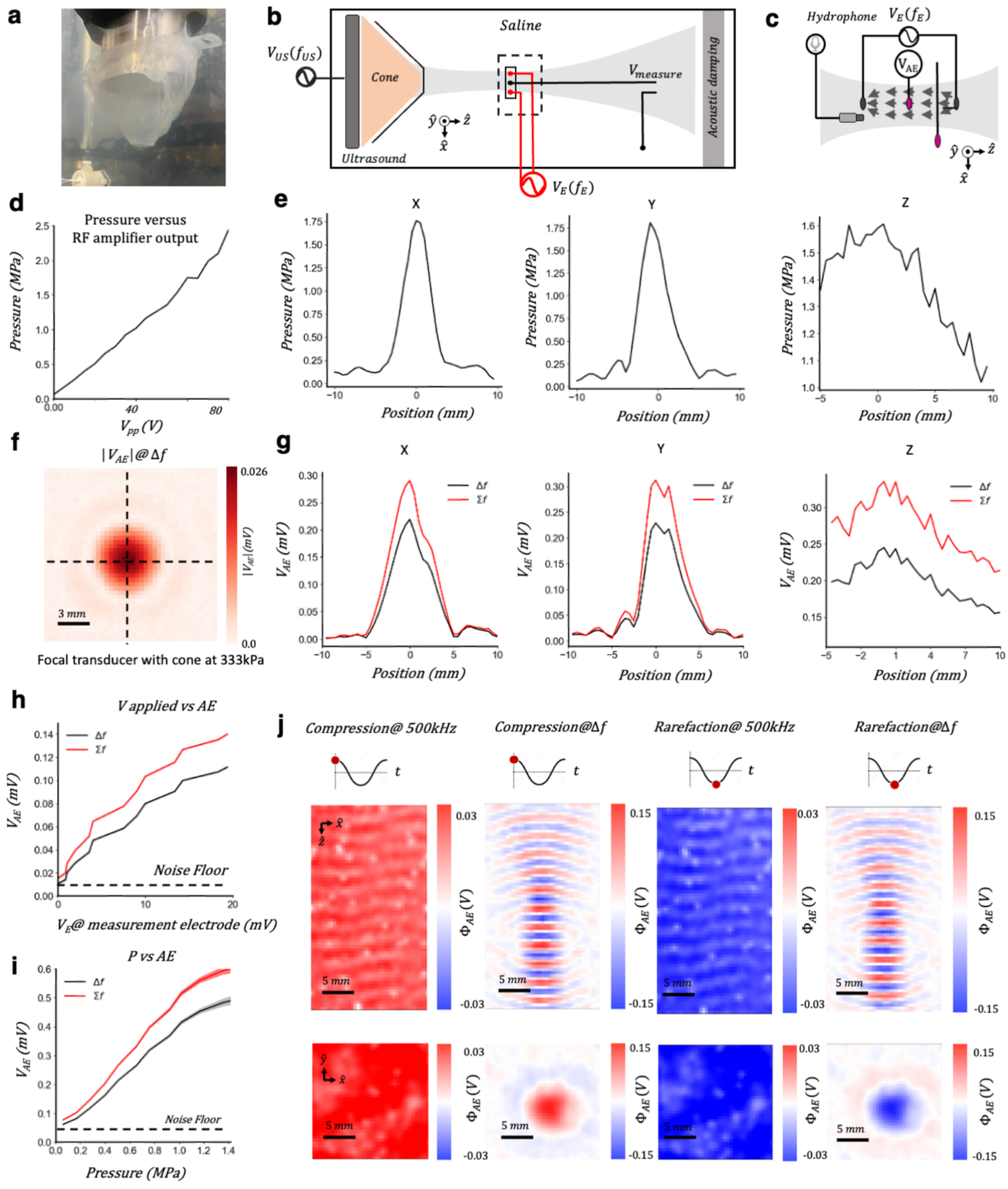


**Figure S1 | Acoustic and acoustoelectric cone characterization a,** 500kHz ultrasound transducer with cone (70mm diameter at the top, 30mm at the bottom, 50mm height), covered in parafilm. **b,** Phantom tank for XYZ scanning of electric fields generated through the acoustic focal area. **c,** hydrophone and electrical measurement electrodes for measuring acoustoelectric effect. **d,** Output voltage of the RF amplifier versus output pressure measured at the focal spot of the ultrasound transducer in 0.9% saline with hydrophone. **e,** XYZ mapping of the acoustic amplitude measured with a hydrophone. Z depth axis is made irregular by reflections off the back of the tank, despite the use of acoustic damping material. **f,** XY spatial distribution map of acoustoelectric difference frequency generated with 500kHz pressure at 333kPa, and 20Vpp@8kHz on voltage electrodes aligned with the ultrasound. 0.1s duration at 5MHz sampling rate, for each pixel. Data band filtered around difference frequency of 492kHz. **g,** Similar parameters to f, except showing X, Y and Z axis sum (508kHz) and difference (492kHz) frequencies and their distribution for each axis. **h,** Ramping amplitude of applied

8kHz voltage with 500kHz pressure constant amplitude at 0.4MPa. Shown is sum and difference voltage amplitudes calibrated to the focal point of the transducer. 8kHz was chosen, so that a shorter time length of recording could be used in each measurement compared to an electric field at a similar frequency to the acoustic signal. **i,** Ramping pressure of 500kHz ultrasound transducer with constant 12V output 8kHz voltage on electrodes. Shown are the sum and difference voltage amplitudes calibrated to the focal point of the transducer. **j,** XY and XZ spatial maps of acoustoelectric difference frequency and acoustic carrier frequency for both a compression and rarefaction.

### 2. F21 material characterization

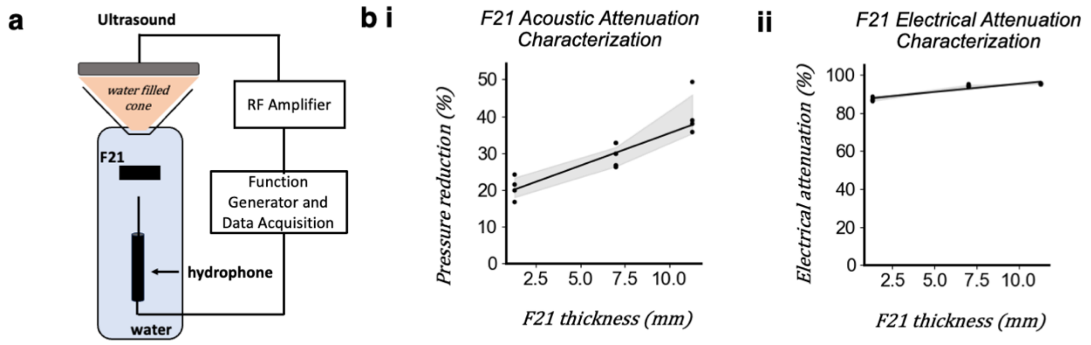


**Figure S2 | F21 acoustic and electrical characterization a,** Hydrophone and transducer arrangement for F21 characterization. **b**, i) F21 material acoustic attenuation versus thickness. 8 second duration recordings were chosen to mimic the continuous field applied in experiments, 5MHz sampling rate repeated 4 times at each F21 thickness. Regression line: Attenuation (%) = 1.75 * thickness (mm) + 17.8 ii) F21 material electrical attenuation versus thickness. Regression line: Attenuation (%) = 0.86*F21 thickness (mm) + 86.

### 3. Impact of changing anaesthesia depth

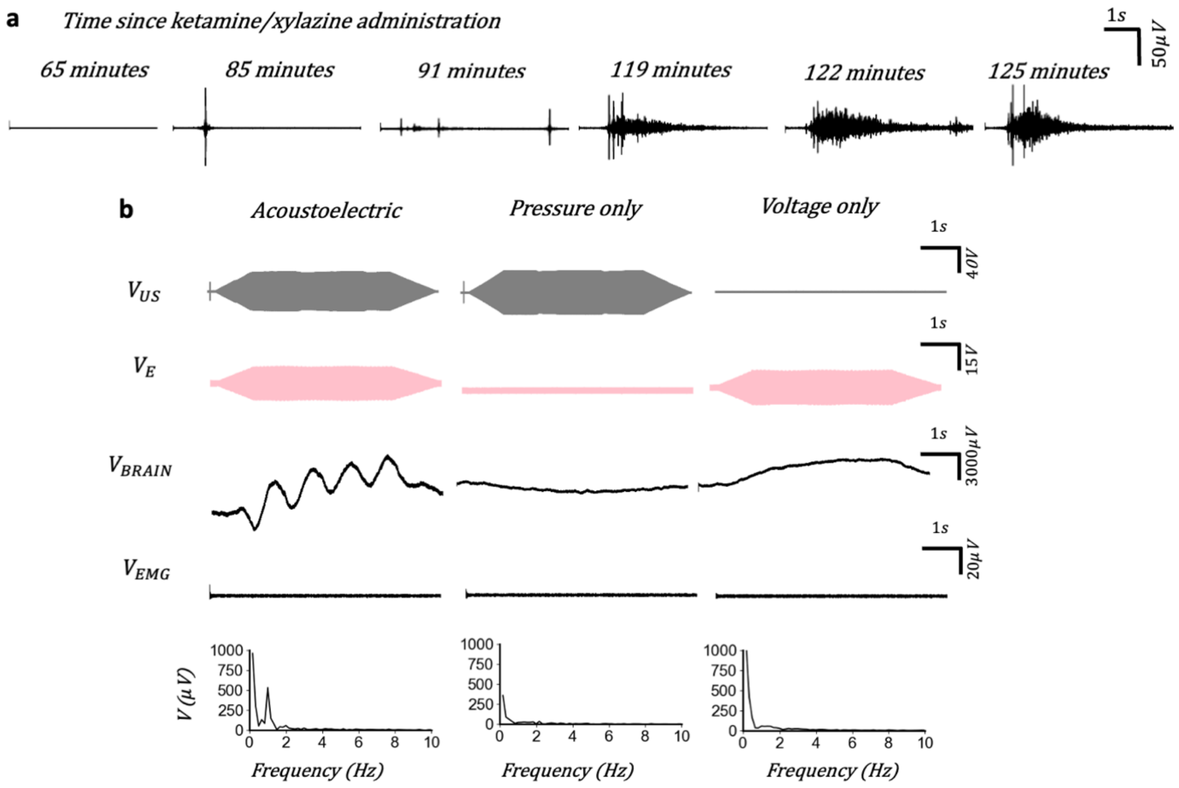


**Figure S3 | Impact of changing anaesthesia depth. a,** Time series showing EMG trace change when ultrasound stimulation is applied at increasing time intervals since the beginning of ketamine/xylazine administration. EMG filtered between 100-1kHz, acoustic signal kept constant in all representative plots (2MPa, 500kHz), sampling rate 5MHz, duration 10s. **b,** Data exclusion example due to depth of anaesthesia; $\Delta f$=1Hz, P=2MPa, V = 30Vpp, Brain signal amplitudes: acoustoelectric signal = $3.396 mV\ pp$, pressure only = $1 mV\ pp$, voltage only = $3 mV\ pp$**.** Leave one out experiment example performed when anaesthesia was too deep to evoke any EMG response, despite Δf=1hz being present at sufficiently high amplitudes in the brain.

We observed large EMG amplitude changes as the anaesthesia level lightened while the applied pressure remained constant (**Fig S3 a**). After administration of anaesthesia, no EMG signal was detected. As the depth of anaesthesia lightened, between 45 minutes to 1hr 15 minutes into the experiment, the EMG response steadily increased. The experimental data collection was performed within the window of lightening anaesthesia where EMG responses could be evoked, and before spontaneous movements began. Once the mouse started moving spontaneously, the experiment was ended. The variance in the EMG response amplitudes due to anaesthesia depth has been discussed in other works[1]. Data were excluded from statistical analyses when the anaesthesia was too deep to evoke an EMG response with example shown in **Fig S3 b**.

### 4. Standing waves in the mouse head

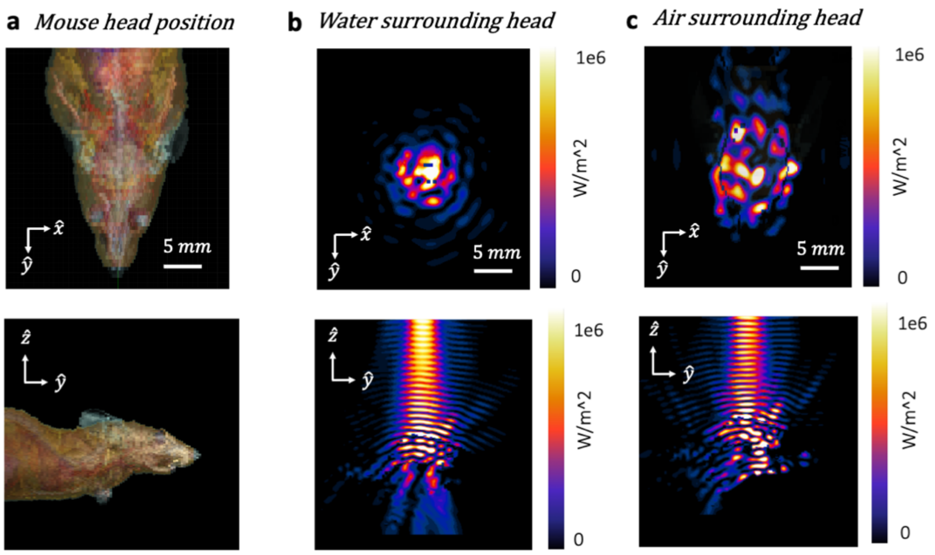


**Figure S4 | Acoustic standing wave reflections. a,** Geometric inhomogeneous mouse model used in Sim4Life finite element modelling simulations with 500kHz ultrasound transducer matching our physical transducer experiments. **b,** 100 acoustic cycles simulated geometry in a mouse surrounded by water. The focal area is clearly in the middle. **c,** 100 acoustic cycles simulated geometry in a with mouse surrounded by air. The acoustic impedance boundary induces emergent standing waves, diminishing the achievable focality at these length scales.

Using the finite element modelling software Sim4Life, we modelled an acoustic wave propagating through an inhomogeneous mouse model with similar ultrasound transducer parameters as our physical experiments (**Fig S4 a**). When the mouse model was surrounded by water the focal area was clear (**Fig S4 b**). However, when the mouse was surrounded by air (**Fig S4 c**), emergent standing waves can be seen. In the case of the mouse, the head size ($\approx 15mm\ diameter$) is smaller than the absorption length in soft tissue of the 500kHz acoustic wave ($\approx 4mm$), diminishing the achievable focality at these length scales. Using either a shorter wavelength transducer or a larger mammal would help mitigate this issue as the absorption length would be decreased.

### 5. Example of poor data included in the statistics: Electrochemical effects at high voltages

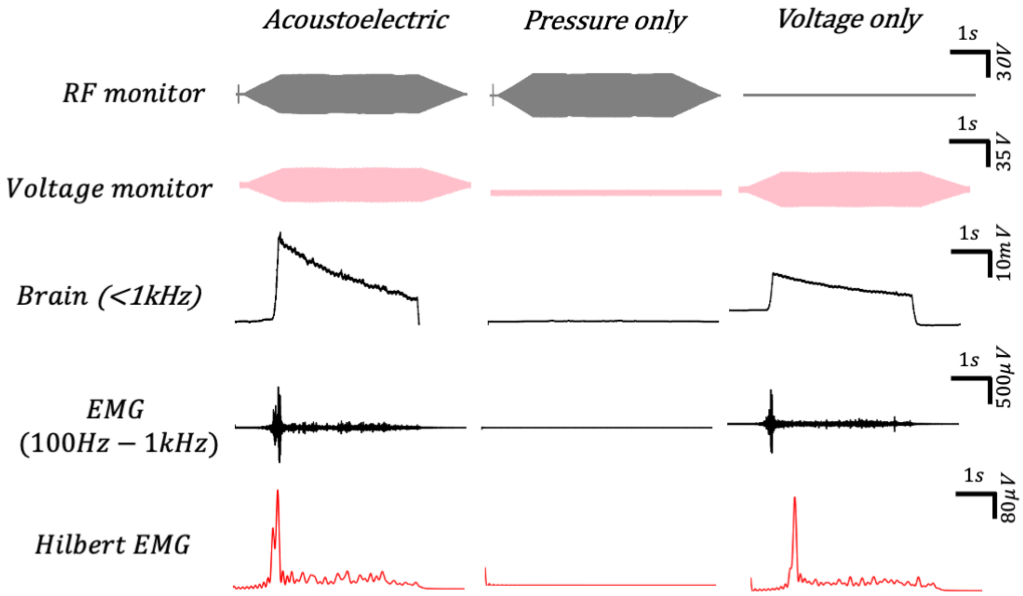


**Figure S5 | Poor data: electrochemical DC offset effects at high voltages.** $\Delta f = 1hz$ P = 1.5MPa, V = 70Vpp. Brain signal amplitudes: acoustoelectric signal = 22.15$mV\ pp$, pressure only = 1.5$mV\ pp$, voltage only = 19.9$mV\ pp$. At very high voltages non-linear electrochemical effects at the electrode interface obfuscate the acoustoelectric difference frequency in the brain signal, presenting as a large DC offset due to the charge imbalance between the stimulation electrodes.

### 6. Acoustoelectric position calibration technique

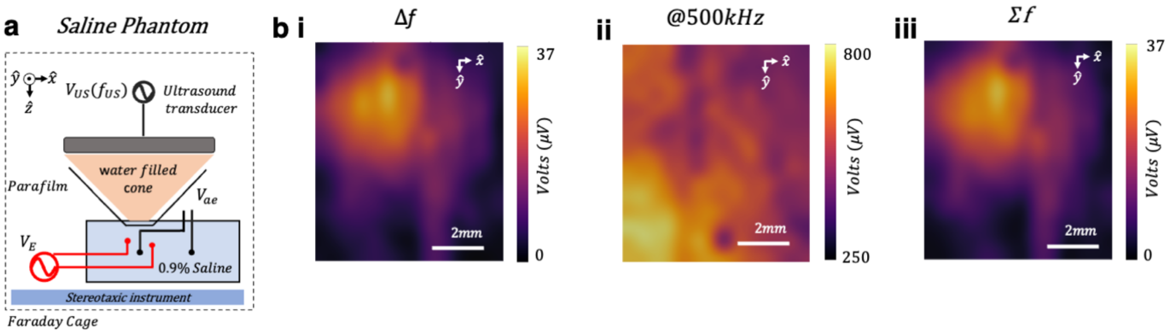


**Figure S6 | Acoustoelectric position calibration. a,** Stimulation and measurement arrangement for position calibration in a saline phantom Note: this method didn't work well in a mouse due to more complex reflections and standing waves and limited experimental timeline. **b,** i) Spatial calibration of stimulation electrode located at visual cortex with 1MPa 500kHz acoustic wave, and 1V output at 8kHz electrical signal on stimulation electrodes directly from function generator, using manual stereotaxic movement to recreate spatial image. i) Δf amplitudes over a 1cm square showing maxima at focus location. ii) Carrier frequency amplitude over same space has no focality. iii) $\Sigma$f amplitudes over a 1cm square showing maxima at focus location.

### 7. Electric frequency mixing does not occur in the signal generation path

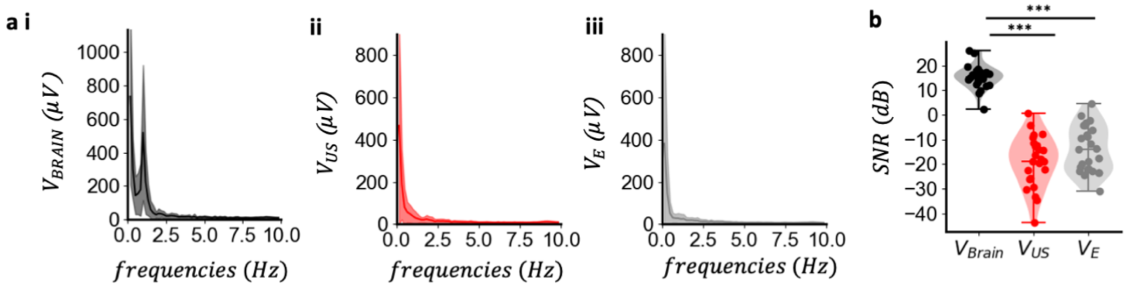


**Figure S7 | In vivo signal generation frequency mixing test. a, i)** *In vivo* measurement of $V_{Brain}$ ASD over the same 1Hz in vivo acoustoelectric data set shown in **Fig 4. ii)** The monitored output from the RF amplifier $V_{US}$ ASD over the same 1Hz in vivo acoustoelectric data as in i) ii) The monitored output from the voltage stimulus $V_E$ ASD is in ii) **b,** Signal-to-noise ratio computed with signal at 1Hz with noise defined as the mean amplitude between 0-2Hz. $V_{Brain}$ is consistently above 0dB, while $V_{US}$ and $V_E$ < 0dB. t-test comparison between SNRs $V_{Brain}$ and $V_{US}$ data at 1Hz frequency, two sided $t_{(21)}$= 15, p = 9.5e-20; t-test comparison between SNRs of $V_{Brain}$ and $V_E$ data at 1Hz frequency, two sided $t_{(21)}$= 14, p = 1.3e-18; $V_{Brain}$ (mean±s.d. = 15.37±4.7 μV); $V_{US}$ (mean±s.d. = -18.81±10.11μV); $V_E$ (mean±s.d. = -14.10±9.16 μV);

The difference frequency could originate at the function generators or recording instrumentation if mixing was to occur in the signal generation path. To show this wasn't the case, we monitored the output stimulation signals for the pressure signal and electric stimulation during the *in vivo* 1Hz neuromodulation experiments reported above. Each stimulation source was generated by a physically separated function generator, each with isolation transformers blocking low frequency signals reflecting from the medium back into the signal generation path. Using the same 1Hz *in vivo* acoustoelectric experiment as shown in Figure 4, the ASD of the measured brain signal, applied pressure and voltage signals was computed showing both mean and s.d. from 0-10Hz (**Fig S7 a i, ii, iii**), such that the 1Hz difference frequency is only visible in the acoustoelectric brain data. To further confirm that electric frequency mixing was not occurring in the stimulus hardware, the signal-to-noise ratio was computed for each of $V_{Brain}$, $V_{US}$, $V_E$ (**Fig S7 b**) where only the voltage measured in the brain has a positive SNR, with significant differences between the brain and output stimulus (p > 0.0005).

### 8. Identifying the source of the RF electrical artefact at 500kHz

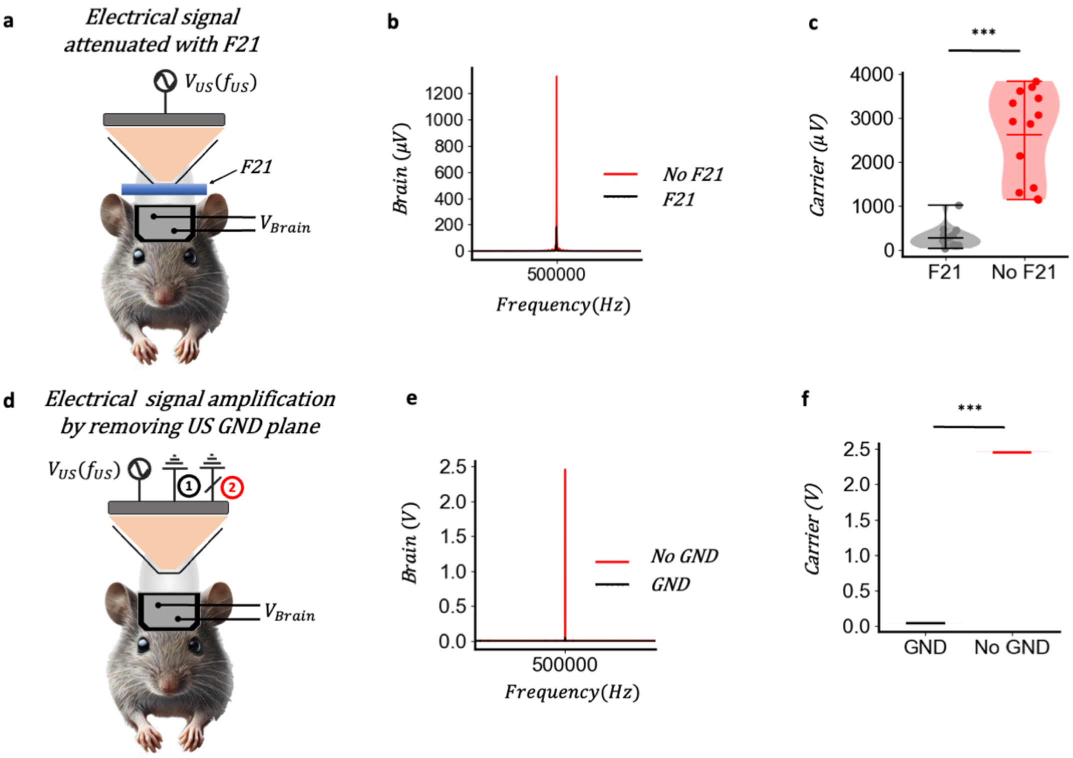


**Figure S8 | Identifying the source of the electrical artefact. a,** *In vivo* experiment arrangement for electrically attenuation and acoustic transparency with 2mm thick F21. **b,** Representative acoustic carrier amplitude ASD comparison between F21/no F21 is placed between the transducer cone and the mouse. **c,** Carrier amplitude comparison, where each mouse was tested with and without F21, with each recording of 6 second duration, using a total of 17 trials over 4 mice. t-test, two sided $t_{(16)}$= -4.99, P = 1.86e-5; F21 group (mean±s.d. = $2003.26 \pm 1410.04\mu V$); without F21 group (mean±s.d. = $270.45 \pm 219.23\ \mu V$). **d,** *In vivo* electric signal amplification experiment removing the ground plane of the ultrasound transducer using a switch installed into the cable. **e,** Representative ASD of large voltage 500kHz artefact when ground plane is removed. **f,** Carrier amplitude comparison with GND connected and disconnected; t-test, two sided $t_{(4)}$= -1207.06, P = 1.47e-22; No GND group (mean±s.d. = $2.46 \pm 3.90 mV$); GND group (mean±s.d. = $0.04 \pm 0.40\ mV$).

In all experiments an electric artefact was measured at the acoustic frequency, despite the separation of the water filled ultrasound cone from the subject through non-conductive parafilm. To determine the origin of this electric artefact, an electrically insulating and acoustically transparent material called F21 was used to attenuate the electric field (Precision acoustics, UK). A 500kHz sinusoid, with a 1MPa acoustic maxima and continuous signal was applied, and the electric signals measured when 2mm thick F21 was inserted between the mouse and the cone (**Fig S8 a, b, c**). There was a difference in the carrier amplitude when this electric shielding test was repeated over 17, 6 second trials (P = 1.86e-5). Conversely, removing the ground connection of the ultrasound transducer, by adding a switch on the transducer cable to disconnected ground while sending through the same 1MPa continuous signal, amplified the electrical transmission measured in the mouse brain as more of the energy is transmitted through the air and into the environment when the ground plane was not available to the transducer as a circuit return (**Fig S8 d, e, f**). Capacitive coupling enabled a high frequency electric field to be transmitted[2].

### 9. Ultrasound induces an electric field at the sum and difference frequency

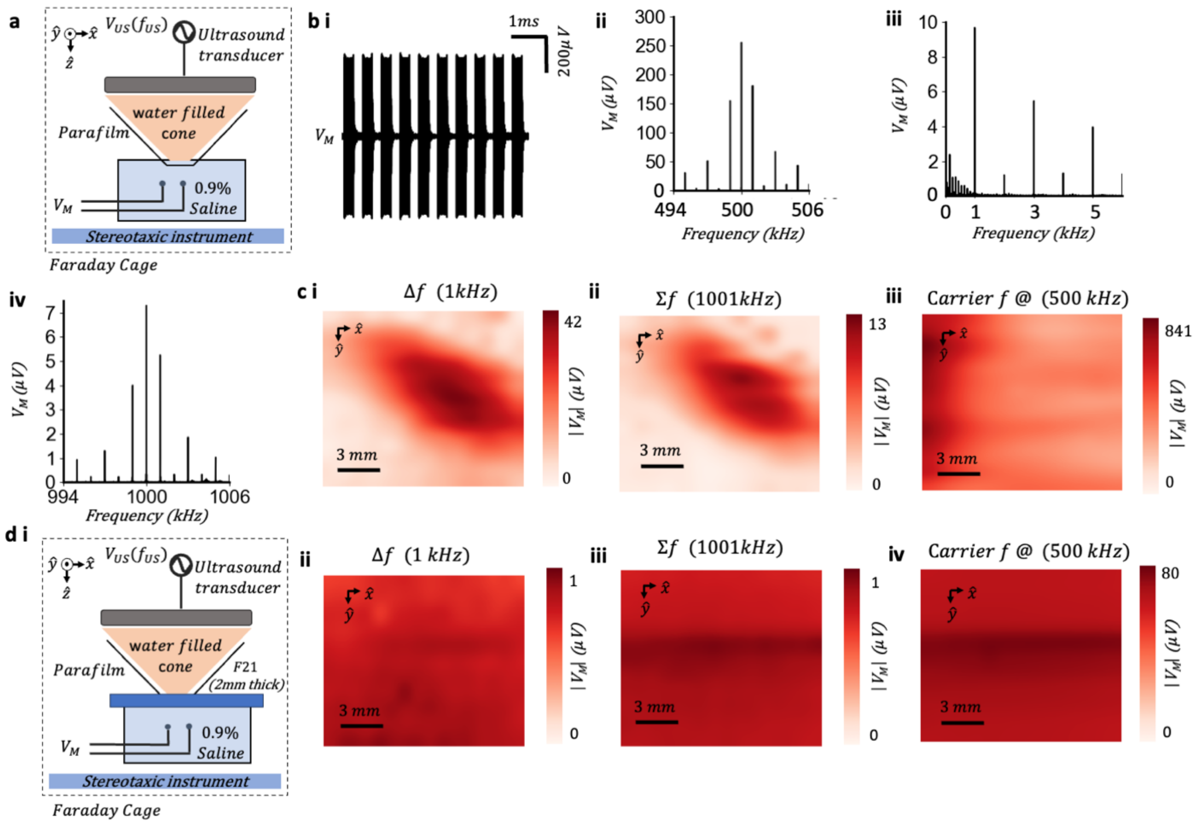


**Figure S9 | Ultrasound undergoes an acoustoelectric interaction induced by the transducer electrical artefact. a,** *in vitro* phantom experimental arrangement in petri dish with 0.9% saline with PRF = 1kHz, carrier frequency at 500kHz, pressure maxima at 1MPa, and stereotaxic instrument moved in 0.5mm increments to produce a spatial map. **b, i)** Representative time series measurement at $V_M$ showing 1kHz PRF (50% duty cycle), with platinum iridium electrodes spaced 7mm apart. **ii)** ASD of the waveform shown in **i)**. **ii)** low frequency ASD around $\Delta f$. **iv)** high frequency ASD around $\Sigma$f. **c, i)** $\Delta f$ spatial map **iii)** carrier frequency map **iv)** $\Sigma$f frequency map. **d, i)** Phantom measurements as in c, with 2mm thick F21 acoustically transparent and electrically isolating material between saline petri dish and end of transducer cone. **ii)** difference frequency shows no electrical focal volume. **iii)** sum frequency shows no focal electrical volume **iv)**. carrier frequency.

We applied only the ultrasound in a saline phantom at a pulse repetition frequency (PRF) of 1kHz and 50% duty cycle (i.e., pulse width 0.5ms) commonly used in tFUS[3] (**Fig. S9 a**). Platinum-Iridium electrodes were placed below the focal volume spaced 7mm apart (**Fig. S9 a,** with instrumentation details in **Methods**). The transducer created an electrical signal in saline at the applied acoustic frequency (**Fig. S9 b i**), showing the original signal spectral peaks decreasing in amplitude around the 500kHz carrier frequency (**Fig. S9 b ii**)**.** When the 1kHz PRF signal multiplies with itself it's modulated to baseband (0Hz), such that the sequentially decreasing amplitude seen around 500kHz now appears around 0Hz (**Fig. S9 b iii**). This decrease in amplitude at baseband, can only be present if the frequency mixing occurred, as echoed by software simulation (**Supplemental Note 10**). Then, at smaller amplitudes the sum frequencies were also present (**Fig. S9 c iii**) alongside further intermodulation products[4].

Next, we investigated the origin of the sum and difference frequencies. We used a stereotaxic instrument to move the ultrasound in 0.5mm increments through an 8mm square, measuring the amplitude of the emergent sum and difference frequencies in the $\hat{x}\hat{y}$ plane (**Fig. S9 c i, ii**), and the applied carrier signal ($f_{US}$) (**Fig. S9 c iii**). The sum and difference frequencies were focal with the acoustic field while the carrier frequency was not focal, matching previous acoustoelectric characterization measurements where an electric field was applied independently at a second frequency to evoke a difference frequency (**Fig. S4 g, j**). Since a sum frequency exists, this focal electric field cannot be induced by the acoustic radiation force (ARF)[5] which only occurs at the low frequency amplitude envelope ($\Delta f$) of the acoustic wave and has not been reported previously as an electrical signal. The focal sum and difference envelope is also not able to be produced through electrical interactions alone, as these would not evoke an electric field focal with the acoustic field.

We then used the acoustically transparent and electrically insulating F21 material to determine if shielding the ionic medium from the electric field from the transducer would decrease the focal mixing at the sum and difference frequencies (**Fig. S9 d i**). The focal area at the sum and difference frequency was greatly attenuated when F21 was in place (**Fig. S9 d ii, iii**), and the electric carrier artefact from the ultrasound was diminished (**Fig. S9 d iv**). This is evidence that the focal electric field is dependent on the electric output of the ultrasound transducer and the focality of the propagating acoustic field i.e. an acoustoelectrically generated field. Similarly in **Fig. 5 d-g** we have shown that when the electric field is independent, the difference frequency is not dependent on the electrical field artefact from the transducer to eliminate electric field only mixing. Hence, we have a multiplication of electric and acoustic fields, which could possibly be explained the acoustoelectric heterodyning effect[6].

### 10. Simulation of $V_{US}$ compared with $V_{US}$ x $P_{US}$

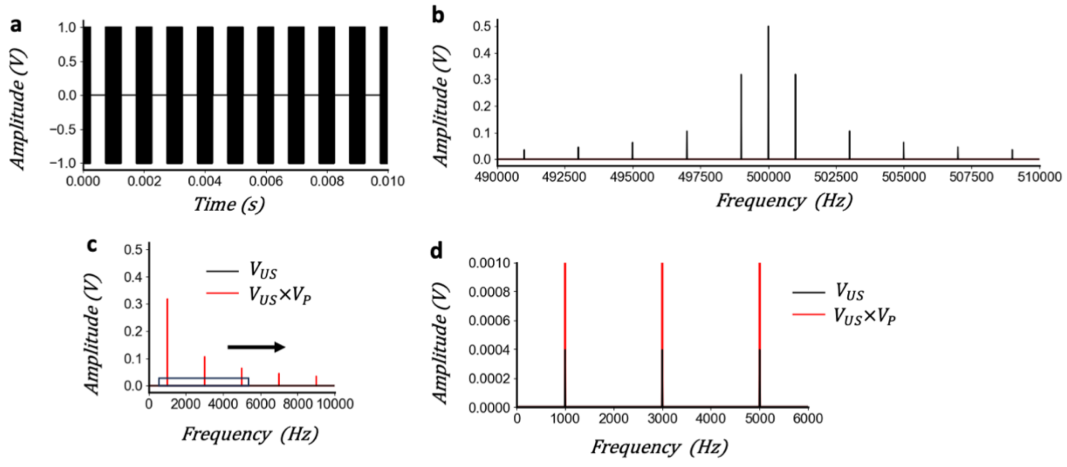


**Figure S10 | Simulation of $V_{US}$ compared with $V_{US}$ x $P_{US}$. a,** $V_{US}$ simulated 500kHz sinusoid with amplitude 1, pulsed at 1kHz. **b,** ASD of $V_{US}$ around the carrier. **c,** ASD at low frequencies comparing $V_{US}$ and $V_{US}$ x $P_{US}$ signal. **d,** Zoom view around the first 3 frequencies, showing residue small $V_{US}$ signals all at similar amplitudes and larger $V_{US}$ x $P_{US}$ signals decreasing in amplitude, mirroring the $V_{US}$ spectrum shape except base banded around 0hz.

### 11. Acoustoelectric Neuromodulation Instrumentation

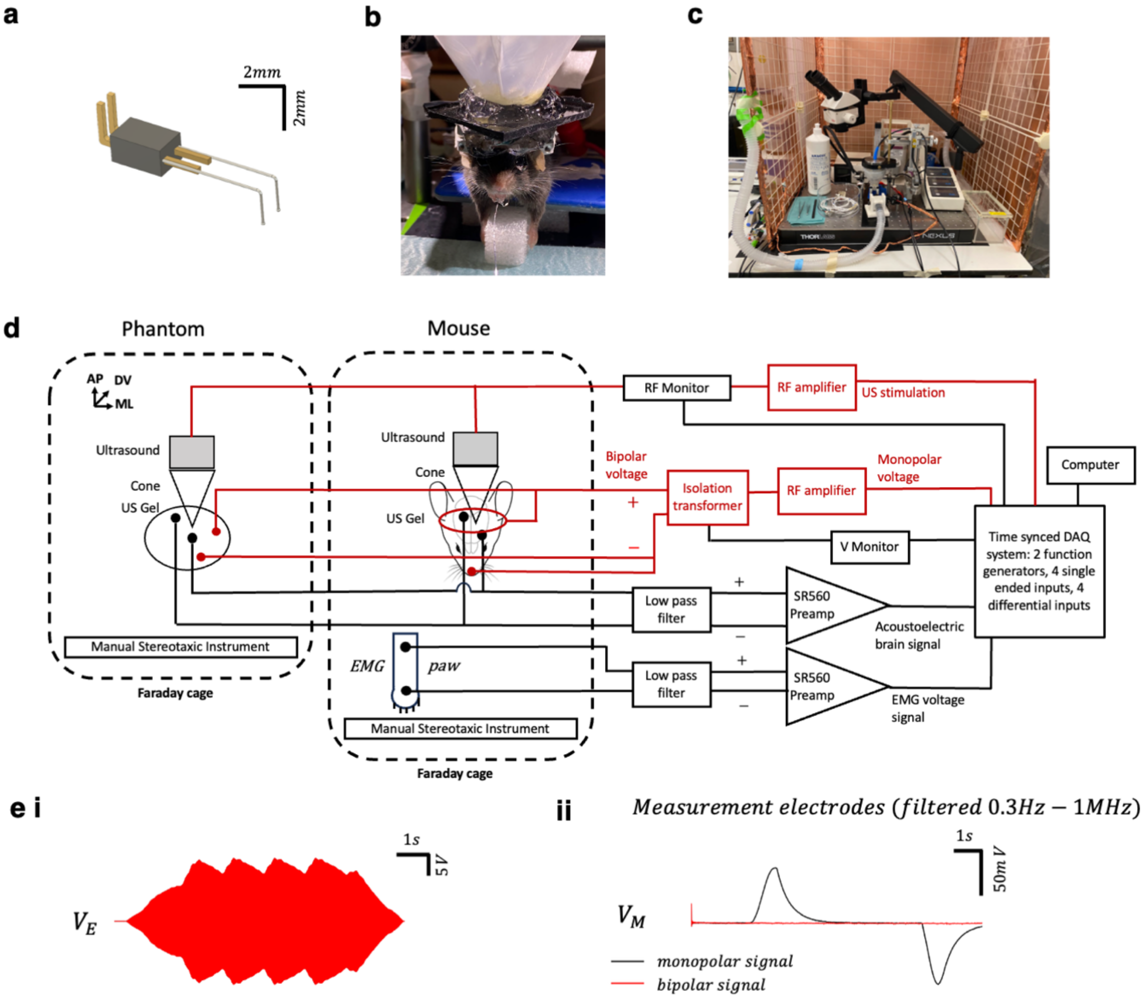


**Figure S11 | Acoustoelectric instrumentation. a,** Electrodes implant design **b,** Mouse with ultrasound cone, gel and 2mm F21 in place. **c,** Experimental apparatus. **d,** Acoustoelectric neuromodulation instrumentation for acoustoelectric neuromodulation in both saline phantom and mouse. Red lines show stimulation equipment and black lines show measurement system. **e,** i) Stimulation signal monitored during 70V peak-to-peak 500kHz voltage signal applied via the Platinum Iridium electrodes used for stimulation in the mouse, some reverberation is present due to electric reflections. ii) Differential measurement at electrodes in 0.9% physiological saline phantom, sampling rate 5Mhz, with differential preamplifier front end filter 0.3Hz-1MHz, gain=1, 12 seconds duration showing monopolar arrangement DC offset at onset and offset (black line) and bipolar signal (red line) where both have been 40Hz software low-pass filtered to reveal the electrochemical DC drift induced between the stimulation electrodes. Note that the preamplifier front end filter was implemented (0.3Hz high pass) to keep the recorded signal within its saturation limits, meaning the recorded filtered signal shows the onset of the DC but not the actual offset.

### 12. Low pass filter

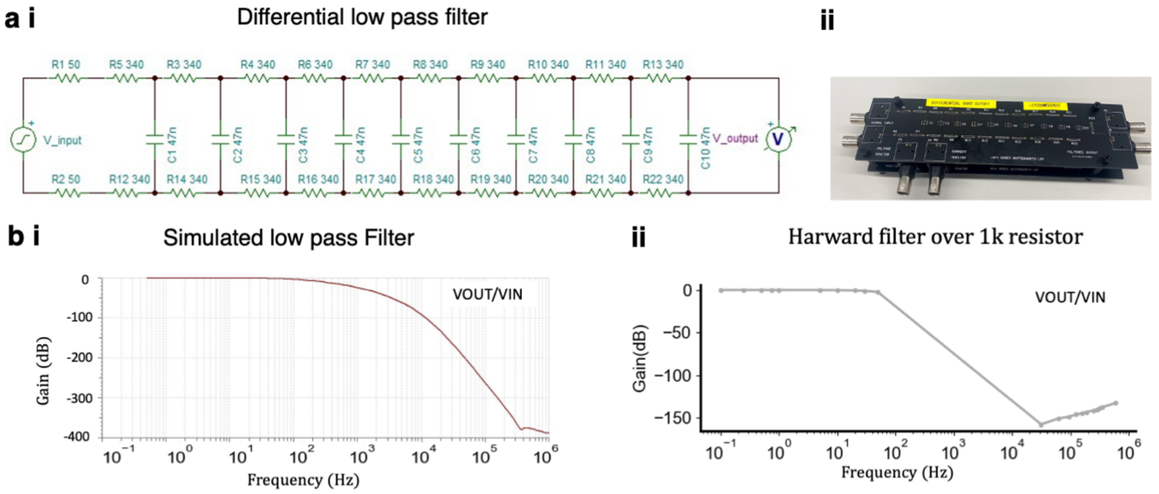


**Figure S12 | Low pass filter a,** i) Circuit for low pass differential filter attenuating 300dB at 500kHz (based on pSPICE simulation). ii) low pass filter was required to filter out the large amplitude high frequency electric signal delivered around 500kHz. **b,** i) simulated transfer function up to 1MHz. ii) Real transfer function of the physical hardware filter.
